# Statins attenuate Wnt/β-catenin signaling by targeting SATB family proteins in colorectal cancer

**DOI:** 10.1101/2024.08.23.609189

**Authors:** Sneha Tripathi, Ekta Gupta, Rutika Naik, Satyajeet Khare, Rafeeq Mir, Saarthi Desai, Swati Humane, Subhash Yadav, Munita Bal, Avanish Saklani, Prachi Patil, Siddhesh Kamat, Sanjeev Galande

**Affiliations:** Laboratory of Chromatin Biology & Epigenetics, Indian Institute of Science Education and Research, Pune 411008, India; Department of Biology, Indian Institute of Science Education and Research, Pune 411008, India; Symbiosis School of Biological Sciences (SSBS), Symbiosis International (Deemed University), Lavale, Pune, India; Present address: Centre for Interdisciplinary Research and Innovations, University of Kashmir, Srinagar, India; Tata Memorial Hospital, Homi Bhabha National Institute, Parel, Mumbai, 400012, India; Centre of Excellence in Epigenetics, Department of Life Sciences, Shiv Nadar Institution of Eminence, Delhi-NCR, India

**Keywords:** Colorectal cancer, Statins, SATB1, SATB2, Wnt/β-catenin signaling, repurposed drugs, tumor-suppressive phenotype

## Abstract

Colorectal cancer is the second leading cause of cancer-related deaths worldwide, highlighting the need for improved treatments and advanced molecular research. A recent therapeutic approach focuses on repurposing drugs to target dysregulated pathways involved in tumorigenesis. Among these, statins, commonly known for lowering cholesterol, have attracted attention for their potential anti-cancer properties. Here, we provide direct evidence for the same by assessing the impact of statin treatment on lipid, transcript, and protein levels. Our findings reveal that statins specifically target key components of the Wnt/β-catenin pathway, a major factor in adenoma formation, including the SATB (Special AT-rich Binding protein) family proteins. While SATB1 is recognized as a regulator of tumorigenesis, particularly under Wnt signaling, SATB2 appears to exert an opposing role. We demonstrate that statin treatment reciprocally alters the expression pattern of these proteins. Furthermore, a human clinical trial evaluating statins as an anti-cancer therapy supports the hypothesis that differential expression of SATB proteins is crucial in tumorigenic outcomes. In conclusion, this modulation by statin treatment suggests promising new therapeutic avenues through drug repurposing.

## Introduction

Colorectal cancer (CRC) is the second most prevalent cause of cancer-related deaths globally, underscoring its severity (1). Despite extensive research efforts aimed at improving outcomes, CRC’s low survival rates, frequent relapses, and limited treatment effectiveness underscore the ongoing challenge it poses. Intensive investigation is directed towards unraveling the molecular underpinnings of CRC and developing targeted therapeutic approaches. Nonetheless, the most promising outcomes continue to be associated with early detection and the removal of polyps, rendering chemotherapy less efficacious in advanced stages for achieving a favorable prognosis. Therefore, there is a growing need for combination therapies that address the early stages of CRC. Researchers are increasingly exploring physiological, genetic and epigenetic factors, as well as environmental factors like diet, to emphasize the importance of chemoprevention. The comprehensive exploration of oncogenic and tumor suppressor pathways in CRC opened up possibilities for therapeutic targets in its early stages. Nevertheless, the field remains dynamic, as new insights into CRC progression continue to emerge with each new study.

Lately, a notable approach in cancer therapeutics involves repurposing drugs that are already approved for treating different disorders. These drugs are chosen because their mechanisms of action are well-understood, targeting physiological pathways known to be disrupted in tumorigenesis. The selection of a repurposed drug primarily hinges on the connection between the drug’s intended target and the observed trends in tumor progression. For instance, there is evidence indicating that individuals with hypercholesterolemia are at higher risk of developing colorectal cancer (2-4). Several studies have also suggested a direct correlation between the cholesterol pathway and oncogenic signaling pathways responsible for tumorigenesis (5-8) and metastasis (9,10). Moreover, CRC patients with elevated cholesterol levels have been reported to experience liver metastasis (11-13). The established association between hypercholesterolemia and CRC prognosis positions statins as a promising candidate for repurposing as an anti-cancer drug.

Statin drugs are the most effective treatment for hypercholesterolemia because they inhibit the mevalonate pathway, which is responsible for cholesterol synthesis. Statins function as competitive inhibitors of HMG CoA reductase, an enzyme that catalyzes the rate-limiting crucial step in the mevalonate pathway, effectively halting the cascade at an earlier stage (14). By doing so, they enable cells to uptake free cholesterol from the bloodstream to fulfill their metabolic requirements, ultimately reducing the blood cholesterol levels. Beyond their pivotal role in managing dyslipidemia, reports have indicated that statins may possess anti-inflammatory properties (15, 16) and can be used to treat coronary heart disease (17). Some studies have explored their anti-neoplastic effects, suggesting that statins can induce apoptosis in breast cancer by targeting mutant p53 (18). A few meta-analyses and early patient cohort studies have shown a positive correlation between statin use and reduced risk of developing CRC (19-21). However, it remains unclear whether this effect is related to or independent of the established mode of action on the mevalonate pathway. Therefore, in our study we have collectively examined the lipid, transcriptome, and proteome profiles in CRC lines upon statin treatment, aiming to establish a mechanistic connection.

Our findings strongly suggest that statins effectively impede the progression of colorectal tumor both in cell cultures and in live animals. The transcriptome and proteome data generated from our statin treatment experiments indicate the emergence of a tumor-suppressive phenotype in CRC cell lines. Additionally, we have observed a specific targeting of the Wnt/β-catenin signaling pathway, which plays a crucial role in the formation of CRC adenomas. Notably, we report a significant decrease in the protein levels of key players within this pathway, including β-catenin, as well as a global regulator, SATB1, following statin treatment.

We focused on understanding the importance of the Wnt/β-catenin canonical pathway in CRC. SATB1 functions as a chromatin organizer and has been shown to interact with β-catenin, leading to the upregulation of Wnt target genes. This interaction creates a feed-forward loop that results in the increased expression of both SATB1 and Wnt target genes (22). Additionally, other independent studies have corroborated these findings, indicating that elevated SATB1 expression is associated with reduced patient survival in CRC (23-25). On the other hand, SATB2, a homolog of SATB1, has generated varying results in the context of CRC. While many reports suggest that SATB2 exerts a tumor suppressive effect (26-30), some studies propose that upregulated SATB2 contributes to tumor progression (31). Despite these observations, there is still no definitive evidence that fully elucidates the distinct roles of SATB1 and SATB2 and their correlation with each other in the development of CRC. We therefore aimed to examine the dynamic expression of SATB proteins, both SATB1 and SATB2, as potential prognostic markers and therapeutic targets, particularly in the context of statin treatment.

Initial investigation revealed that statins have an opposing effect on SATB1 and SATB2 proteins in colorectal cancer (CRC). Specifically, we observed a time- and dose-dependent downregulation of SATB1 in response to statin treatment (32). Here, we further demonstrated that statins effectively reverse the expression patterns of SATB1 and SATB2 in both 2D cell cultures and 3D spheroid model systems, leading to a reduction in tumor burden in *in vivo* experiments. These findings are primarily observed at the protein level and can be rescued by the supplementation of mevalonate in cell culture. Moreover, in line with our laboratory findings, a human study conducted during the phase II/III clinical trial, involving the administration of statins to CRC patients, reinforces the potential of SATB proteins as valuable prognostic and therapeutic targets. Our comprehensive approach, which incorporates multi-omics data and employs various model systems, significantly contributes to our understanding of the molecular mechanisms underlying the anti-tumor effects of statins.

## Results

### Lipid profile of statin treated cells exhibits a tumor suppressive phenotype

We examined the lipid, transcript, and protein profile in colorectal cancer (CRC) cells treated with statins to gain a comprehensive understanding of the physiological and tumorigenic pathways in synergy. Our primary focus was on the lipid profile due to the known effect of statins on the cholesterol biosynthesis pathway. Assessing the cholesterol status in CRC lines during statin treatment was crucial for validating its mode of action. Beyond cholesterol, our targeted lipidomics analysis included the evaluation of oxysterols, prostaglandins, and Free Fatty Acids (FFAs) known for their involvement in tumor progression (33-37).

In our experimental setup using the CRC cell line HCT116, we observed a significant reduction in cholesterol levels upon statin treatment (Figure 1). Subsequently, the levels of oxysterols, cholesterol derivatives, exhibited a notable reduction. In addition to oxysterols, prostaglandins are also essential for signaling cascades, as they play a crucial role in relaying information in metabolic pathways (38-41), cell division and differentiation (42). Therefore, we monitored the levels of prostaglandins upon statin treatment as well, however, we did not observe a significant alteration in the levels of any of the species (Figure 1A).

**Figure 1:**
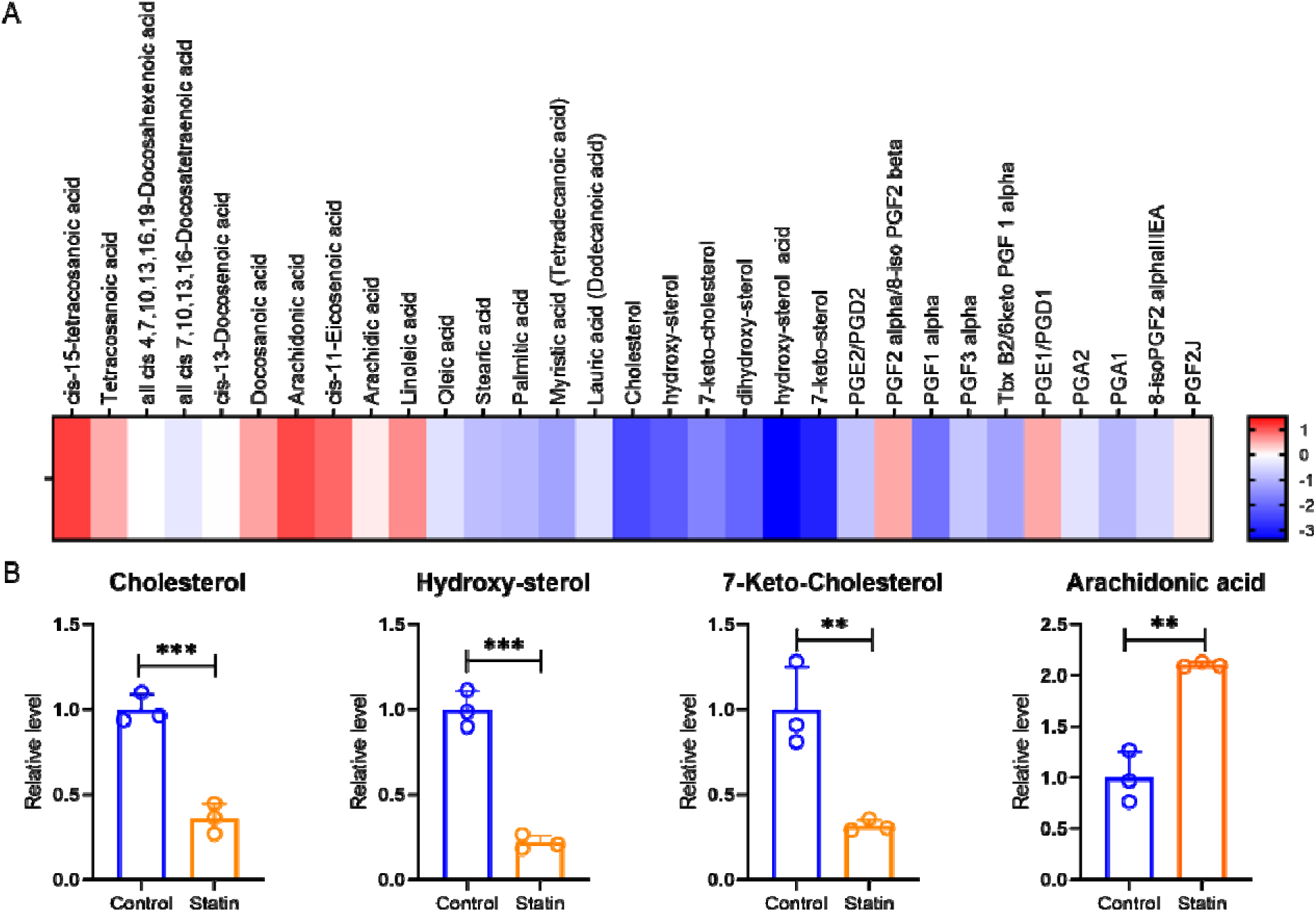
Statin downregulates cholesterol and its derivatives in CRC cells, validating the canonical mode of action. (A) Heatmap to represent the Log 2-fold change levels of Free Fatty Acids (FFAs), Cholesterol, Oxysterols, and Prostaglandins in Statin treated CRC line HCT116. No alteration was observed in levels of Prostaglandin species and most of the FFAs. However, oxysterols were significantly downregulated. (B) Graphs to plot the Relative levels of Cholesterol, Hydroxy-sterol, 7-Keto-Cholesterol, and Arachidonic acid on statin treatment as compared to the untreated control set. Significant reduction was observed in cholesterol and its derivatives; however, arachidonic acid was upregulated. Biological replicates n=3, Students t-test was performed for statistical analysis **p<0.001, ***p<0.0001

Most FFAs species in our analysis remailed unaffected, except for stearic acid, palmitic acid, and myristic acid (Tetradecanoic acid) which showed a significant decrease upon statin treatment (Figure 1A). However, there was a slight increase in arachidonic acid levels, a lipid associated with inflammation (Figure 1B). The inflammatory pathway is regulated by a cascade involving prostaglandins, cytokines, and the NF-κB pathway (43). Although statin treatment is known to reduce inflammation by downregulating cytokines such as IL-6 and IL-8 (44), our transcriptomics analysis did not reveal alterations in prostaglandin levels or other components of the inflammatory pathway. Therefore, the elevated arachidonic acid levels in our lipidomics data might signify a stress response in cells rather than activation of the inflammatory signaling.

Our targeted lipidomics study overall indicated a reduction in cholesterol levels, affirming the intact mode of action of statins in CRC cells. This reduction in cholesterol also led to the downregulation of oxysterols. While prostaglandins and most FFAs remained unaffected, the observed changes, particularly the reduction in stearic acid, palmitic acid, and myristic acid, suggest a potential tumor-suppressive lipid profile associated with statin treatment. This is noteworthy considering the reported surge and accumulation of lipids in rapidly dividing tumor cells, where cholesterol and FFAs play crucial roles in membrane building, immune response modulation, and drug resistance (8, 45-49). The dysregulation of FFAs further contributes to tumorigenesis by influencing tumor progression or microenvironment remodeling (50-53). In summary, the statin mediated reduction in cholesterol may indicate a favorable impact on the lipid profile, aligning with its known anti-tumorigenic effects.

### Transcriptome analysis of Statin treated cells reveals effect on multiple tumorigenic pathways including Wnt/β-catenin signaling

The importance of examining the entire genetic transcript level during statin treatment lies in its effect on genes responsive to the cholesterol biosynthesis pathway. It is established that a decrease in cholesterol levels leads to the upregulation of genes involved in cholesterol biosynthesis. The key transcription factor, SREBP2, is sensitive to cellular cholesterol levels through mevalonate and oxysterols. When cholesterol levels decrease, SREBP2 activates genes in the mevalonate pathway, which then replenish cholesterol through neogenesis or uptake. In our transcriptome study, we initially focused on the effect of statin treatment on cholesterol responsive genes in the HCT116 CRC cell line. As anticipated, the statin mediated reduction in cholesterol resulted in a significant upregulation of mevalonate pathway genes (Figure 2A). Quantitative RT-PCR analysis of HMGR (coding for HMG CoA reductase) and SREBF2 (coding for SREBP2) expression levels supported this observed increase (Figure 2D). Gene Ontology (GO) term analysis reported that the other upregulated set of genes (Figure 2A, Supplementary Table 1) were associated with fatty acid synthesis, miR33 activity, Omega 9 fatty acid synthesis, and Interleukin 2 family signaling. Notably, the predicted upregulation of transcription factors such as KLF15, and AP2 is known to play a tumor-suppressive role (54, 55).

**Figure 2:**
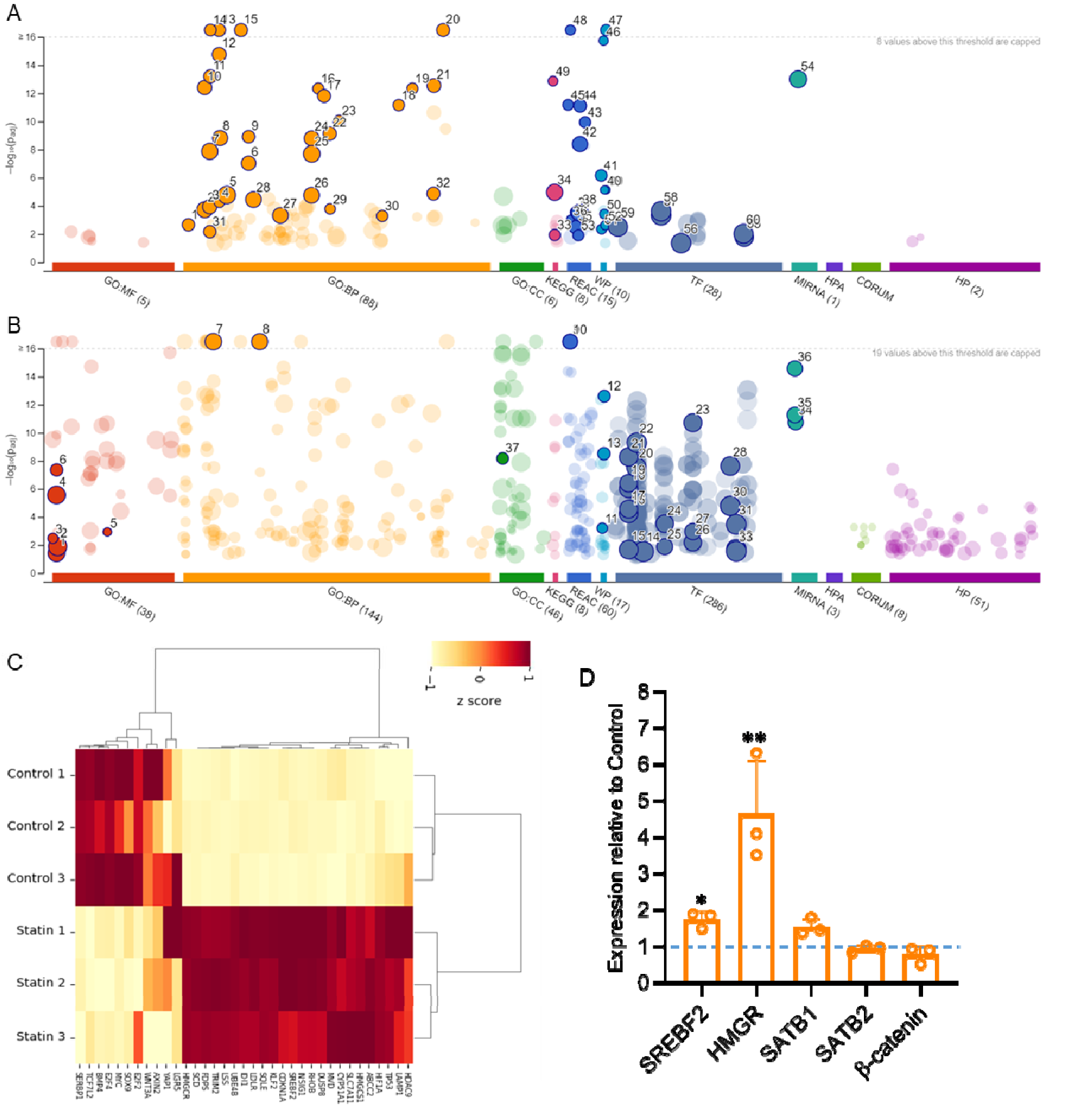
Transcriptome analysis of HCT116 cells upon statin treatment reveals a tumor-suppressive phenotype. Gene Ontology terms were plotted using g:Profiler and observed separately for upregulated (A) and downregulated (B) set of genes on statin treatment in HCT116 cell line. The upregulated GO terms specifically pertain to cholesterol biosynthesis pathway genes, whereas the downregulated terms are related to tumor progressive transcription factors. (C) Heatmap for the DE genes significantly expressed in control versus statin-treated sets. The heatmap was made by selecting a few representative genes of the cholesterol pathway and Wnt target genes using FLASKI online portal. [Iqbal, A., Duitama, C., Metge, F., Rosskopp, D., Boucas, J. Flaski. (2021). doi:10.5281/zenodo.4849515] (D) Validation for transcript level alteration of SREBF2, HMGR, SATB1, SATB2 and β-catenin was performed on statin treated cells. Biological replicates n=3,*p<0.05, **p<0.005 according to Students’ t test analysis.

The GO terms for the downregulated genes were separately plotted (Figure 2B, Supplementary Table 2). Transcription factor (TF) activity prediction indicated an overall reduction in tumorigenic factors. A prominent GO term in this set was the Gastric cancer network, emphasizing the canonical Wnt pathway. Some of the listed transcription factors with reduced activity, including c-Myc, p300, E2F1/2, and TCF-1, are known to be regulated by major molecular players of Wnt/β-catenin signaling (56). We plotted a heatmap to collectively compare the expression profiles of Wnt target genes and cholesterol-responsive genes (Figure 2C). The analysis revealed a distinct downregulation of Wnt-responsive genes and upregulation of cholesterol-responsive genes upon statin treatment.

Given the central role of Wnt/ β-catenin signaling gastric cancer ontology and its pivotal role in the initiation of colorectal cancer (CRC), we investigated whether statin treatment affected downstream effectors of this pathway. The APC and β-catenin mutations predominantly result in aberrant Wnt/ β-catenin signaling, and approximately 75% of CRC patients harbor these mutations (57, 58). Therefore, we reasoned that if statin treatment results in a reduction in downstream effectors of this pathway, then it might target the Wnt/β-catenin signaling via its upstream players. To validate this, we assessed the transcript level of β-catenin, the major Wnt pathway player, and interestingly observed no alteration. Despite no change in the transcript levels of β-catenin, a major Wnt pathway player, as observed in our analysis and confirmed by quantitative RT-PCR analysis (Figure 2D), we further validated the expression of SATB1, a crucial upstream factor with known tumorigenic effects (22, 59). The transcriptome data and quantitative RT-PCR validation showed no alteration in SATB1 transcripts upon statin treatment. Similarly, SATB2, a homolog known for its anti-tumorigenic effects (26), exhibited no change at the transcript level (Figure 2D). Taken together, the transcriptome analysis revealed that statin treatment upregulated cholesterol biosynthesis genes due to a feedback mechanism resulting from lowered cholesterol. Conversely, statin mediated downregulation of oncogenic players targeted Wnt-responsive genes. Intriguingly, major upstream players of the Wnt pathway, including β-catenin, SATB1, and SATB2 did not exhibit changes at the transcript level. Therefore, the observed reduction in the transcript levels of Wnt responsive genes may be attributed to the effect of statin on the protein levels of these upstream molecules.

### Proteome analysis of Statin treated CRC cell line affirms downregulation of Wnt/ β-catenin signaling

To further confirm the effect of statin treatment, we performed whole-cell lysate MS-MS analysis on the HCT15 CRC cell line treated with statin. The proteome data allowed us to establish a connection between transcriptional and translational outcomes, offering a comprehensive view of the affected pathways with direct translational implications. Remarkably, we observed that statin treatment did not significantly affect most proteins, providing additional evidence for the specificity of statin’s mode of action. The commonly categorized off-target effects of the drug, referred to as pleiotropic outcomes, are predominantly linked to its impact on the mevalonate pathway. However, the lack of substantial alterations in the majority of protein levels suggests a tightly regulated mechanism mediated by statin.

Moreover, there was no subset of proteins that exhibited significant upregulation. Interestingly, the set of proteins displaying significant alterations were those undergoing downregulation. Therefore, we performed a GO term analysis for the downregulated proteins to discern the affected pathways (Figure 3A, Supplementary Table 3). The most noteworthy result was the suppression of Wnt/ β-catenin signaling, EGFR signaling, and TGF-β signaling, all of which contribute to tumorigenesis. The prediction of reduced transcription factor activity included E2F1/2, p300, and Sp1. We plotted peptide counts for major players such as β-catenin, YAP, and CUL-3 E3 ubiquitin ligase, known for their roles in the Wnt/ β-catenin pathway (60) (Figure 3C). The relative expression of these proteins was significantly downregulated upon statin treatment, while the expression of housekeeping genes such as actin and GAPDH remained unchanged.

**Figure 3:**
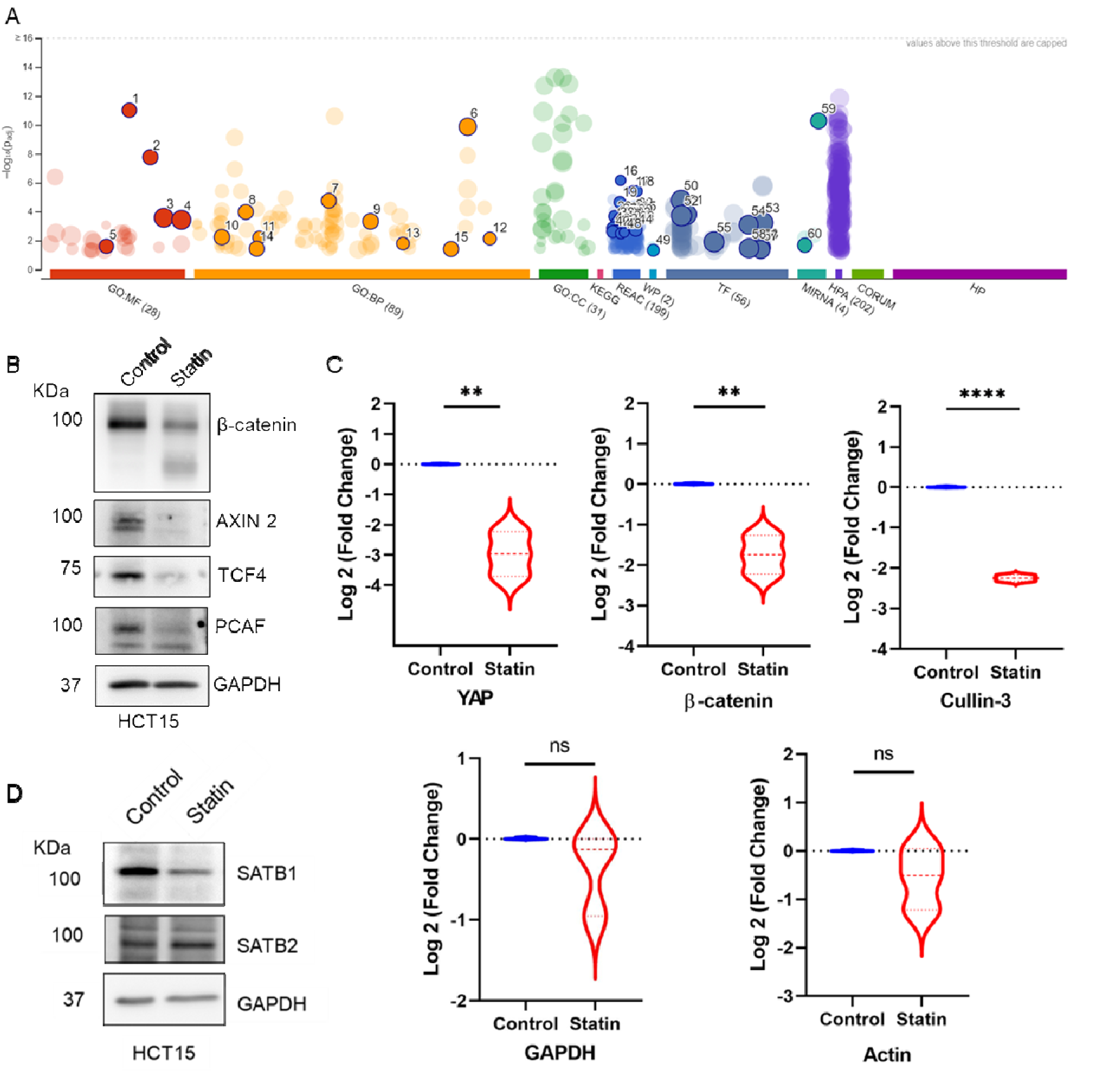
The proteomics profile depicting significantly downregulated proteins upon statin treatment reveals Wnt signaling as a target. (A) Downregulated proteins represented in GO term (plotted on g:Profiler) on statin treatment in HCT15 cell line. (B) Validation of the effect of statin treatment on Wnt target proteins by immunoblotting. A significant reduction at the protein level of Wnt-responsive players is observed on statin treatment. (C) Log fold change was observed in the peptide counts of YAP, β-catenin, and Cullin-3 in control versus statin-treated proteomics analysis, whereas no alteration was observed in housekeeping proteins like GAPDH and Actin. Biological replicates=3 with statistical significance of **p<0.005, ****p<0.00005, ns is non-significant according to Students’ t-test analysis. (D)Immunoblots to delineate SATB1 and SATB2 protein levels in HCT15 cell line upon statin treatment. SATB1 protein was significantly reduced, whereas SATB2 expression slightly increased on statin treatment.

We further confirmed the effect of statin by probing immunoblots for key components of the Wnt pathway, including β-catenin, AXIN 2, TCF4, and PCAF (Figure 3B, Supplementary Figure 1A). Our findings revealed a significant decrease in the levels of all these proteins. Thus, statin treatment led to the downregulation of Wnt-responsive genes at both the protein and the transcript levels. Notably, the upstream regulator β-catenin appeared to be affected only at the protein level, supporting our conclusion that the transcript level changes in the downstream Wnt targets were presumably mediated by the downregulation of the upstream player at the protein level.

Given the known involvement of SATB1 in Wnt/ β-catenin signaling, we probed an immunoblot to assess the levels of SATB1 and its homolog SATB2. Intriguingly, SATB1, which is tumorigenic, exhibited downregulation, while the opposite occurred for SATB2, known to function in contrast to its homolog, showing no significant alteration upon statin treatment (Figure 3D, Supplementary Figure 1B). While previous reports have highlighted the opposing roles of both SATB family proteins, their reciprocal expression in various tissues remains less established. Therefore, a comprehensive understanding of the dynamics of the expression of SATB1 and 2 expression is crucial for considering them as therapeutic targets of statins.

These findings provide insights into the specificity of statin’s anti-tumor mechanism in CRC, particularly targeting the Wnt/ β-catenin signaling. Our proteome data establishes the relevance of the statin-mediated regulation by specifically targeting tumorigenic players.

### Mevalonate supplementation rescues the effect of statin at the protein level

As described previously, statins act on the mevalonate pathway by inhibiting the rate-limiting step catalyzed by HMG CoA reductase. The lactone ring of statin competitively inhibits its substrate, HMG CoA, thus preventing the formation of mevalonate - a precursor essential for cholesterol synthesis. Consequently, we sought to examine the effect of mevalonate supplementation on CRC cell lines when combined with statin treatment.

The presence of mevalonate appeared to counteract the statin-induced effects on the upstream components of the Wnt pathway. This was evident as the protein levels of β-catenin and SATB1 were restored, while the SATB2 levels remained unchanged (Figure 4A, Supplementary Figure 1C). Consistent with earlier findings, the transcript levels of these components remained (Figure 4B, Supplementary Figure 2). The addition of mevalonate, the product of the enzyme inhibited by statins, facilitated the restoration of the downstream pathway, thereby reversing the effects of statin. However, the cells treated with mevalonate and statin exhibited cholesterol levels comparable to the untreated control group, confirming the rescue in the downstream cascade (Figure 4C).

**Figure 4:**
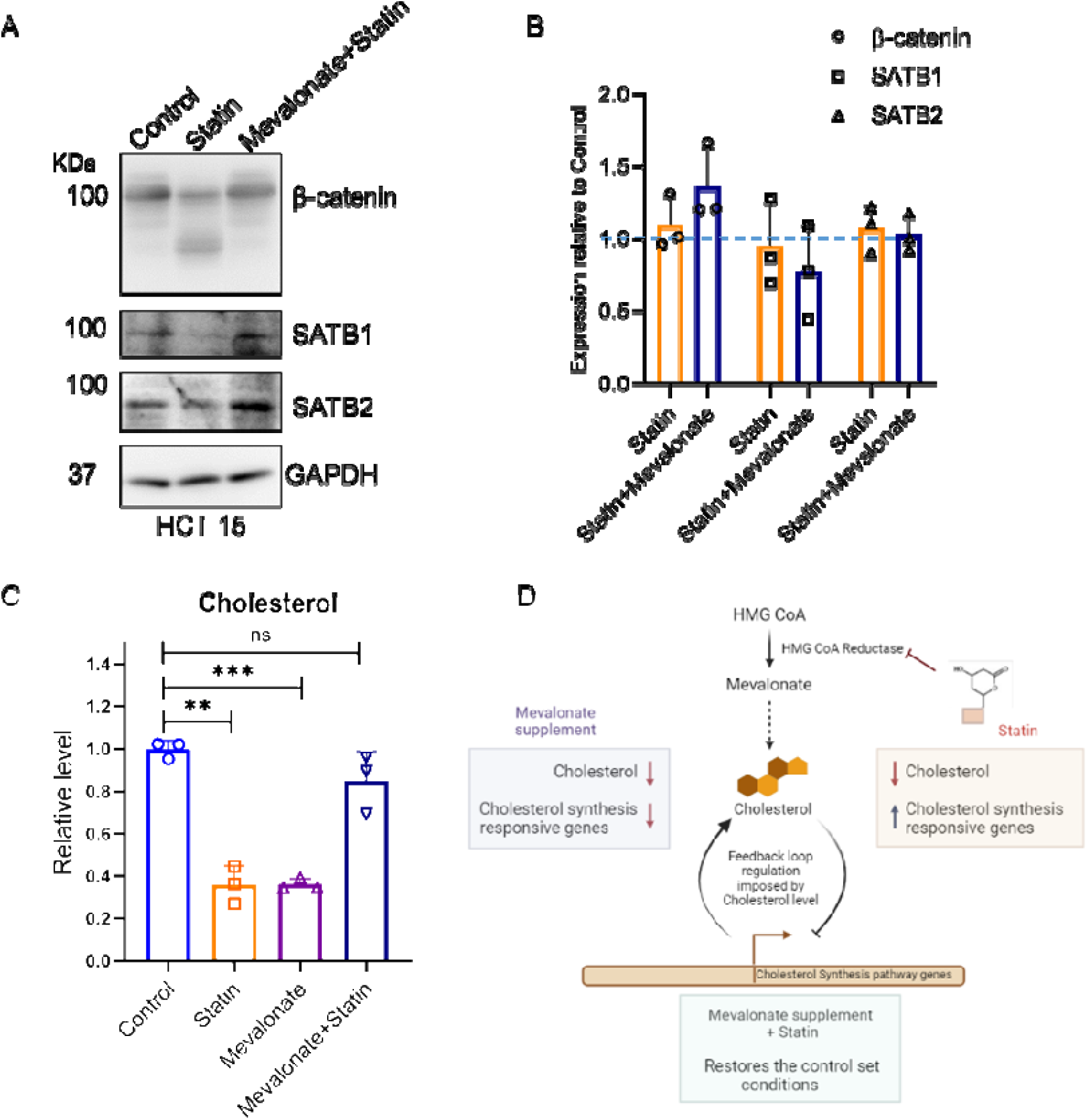
Mevalonate supplementation rescues the effects of statin on SATB proteins as well as on cholesterol levels. (A) Immunoblots for protein level alterations of β-catenin, SATB1 and SATB2 on statin treatment and mevalonate supplement. Rescue in the protein levels were observed on mevalonate supplementation. (B) Relative expression of β-catenin, SATB1 and SATB2 to observe transcript level changes upon statin treatment and mevalonate supplement. No significant alteration was observed in any of the genes. (C, D) Relative levels of cholesterol on mevalonate supplement and statin treatment supported by an illustration depicting the effect of the two conditions on the feedback mechanism at play for the cholesterol-responsive genes. The cholesterol levels were reduced on statin treatment and restored on mevalonate supplement as compared to the untreated control. However, the reduced levels of cholesterol in only mevalonate condition might be a result of the feedback machinery that downregulates cholesterol responsive genes in the presence of excess mevalonate. Biological replicates n=3, **p<0.05, ***p<0.005, ns is non-significant as per Students’ t-test analysis.

Interestingly, the sole set of cells supplemented with mevalonate showed a reduced cholesterol level compared to the untreated control. This peculiar observation may be explained by considering the feedback mechanism that regulates the transcription of the genes responsible for cholesterol biosynthesis in response to the presence of mevalonate (Figure 4D). In this scenario, the surplus mevalonate led to the downregulation of responsive genes, causing the cholesterol synthesis machinery to shut down. We hypothesize that the existing cholesterol might have been utilized by the time of cell harvesting, and with no replenishment from the synthesis apparatus, a decreased cholesterol level was observed in the LC-MS readout.

These results underscore the specificity of statin’s effect on CRC, ruling out the possibility of a generic drug response artifact. Statin unequivocally affects the Wnt/ β-catenin signaling by selectively downregulating key molecular players at the protein level, a phenomenon reversed by mevalonate supplementation. Notably, transcript levels remained unaffected under all conditions. The restoration of statin-mediated effects suggests a regulatory mechanism in Wnt/ β-catenin signaling, potentially linked to the inhibition of the mevalonate pathway.

### 3D spheroids derived from CRC cell lines exhibit distinct expression pattern of epithelial and mesenchymal markers and Wnt players

To further confirm the effect of statins on CRC, we opted for a more intricate model system than conventional two-dimensional monolayers (2D cell cultures). Hence, we established a 3D spheroid culture to investigate colorectal tumorigenesis (Figure 5A). The significance of spheroids has been well recognized in the cancer biology community as a model system superior to 2D cell cultures in accurately representing the cellular and molecular transitions during tumorigenesis (61). However, before delving into statin-mediated regulation, we initially validated the spheroids for oncogenicity by assessing the Epithelial-to-Mesenchymal Transition (EMT) and Mesenchymal-to-Epithelial Transition (MET) markers in comparison to 2D cells.

**Figure 5:**
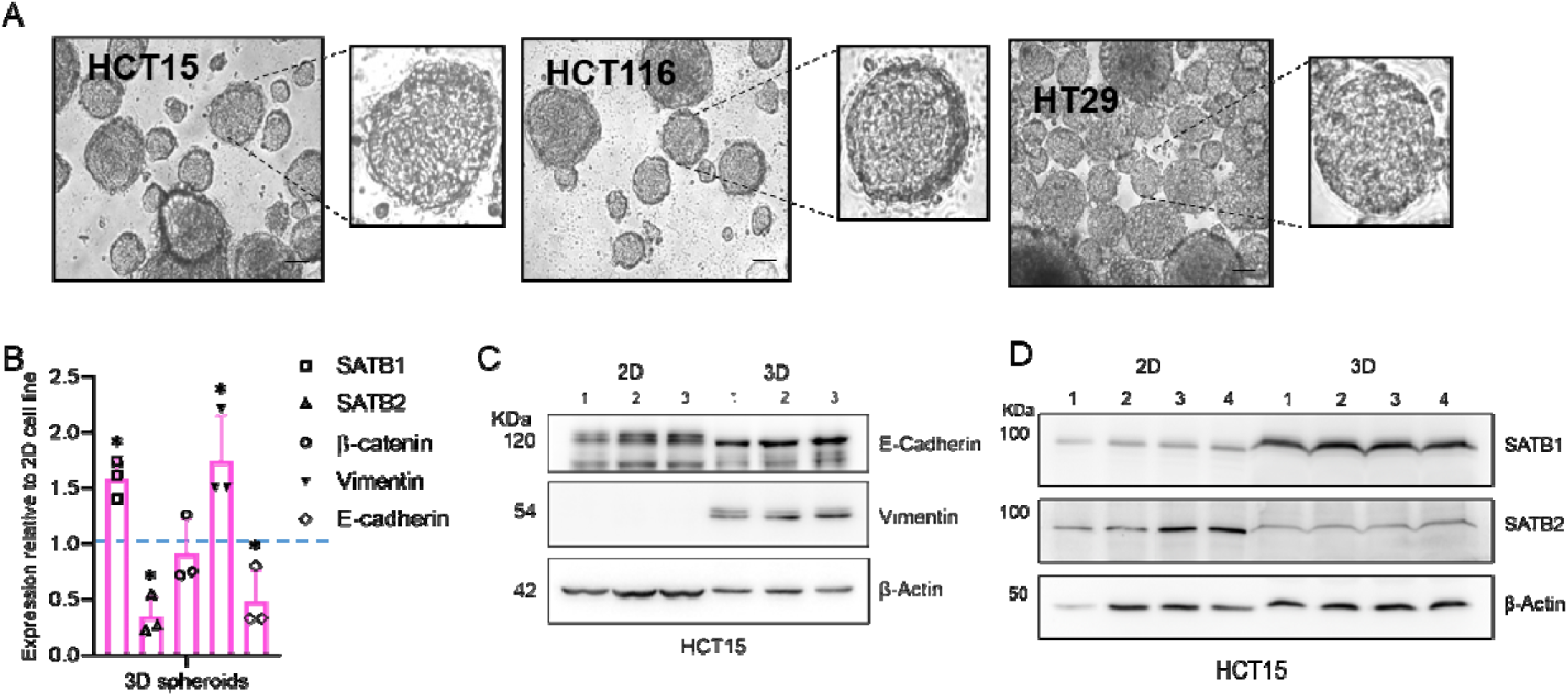
3D spheroids of CRC cells recapitulate the EMT phenotype better than 2D cells. (A) Images of spheroids for CRC lines HCT15, HCT116, HT29 which were allowed to form in matrigel for 4 days. Phase contrast images of spheroids, scale bar 50 μm, with zoomed in image in adjacent box. (B) Relative gene expression of SATB1, SATB2, β-catenin and EMT-MET markers, Vimentin and E-cadherin, respectively. The spheroids showed higher expression of EMT marker Vimentin along with higher expression of SATB1 than SATB2. β-catenin did not show any alteration. (C) Immunoblot to validate the protein levels of the EMT marker Vimentin and MET marker E-cadherin in 2D cell line HCT15 versus the 3D spheroids of HCT15 cells. The protein levels of Vimentin are significantly upregulated in 3D spheroids. (D) Immunoblot to observe the protein levels of SATB1 and 2 in 2D and 3D spheroids of HCT15 cells recapitulating the tumorigenic phenotype of spheroids. Biological replicates n=3, *p< 0.05, **p<0.005 as per Students’ t-test analysis.

The 3D spheroids exhibited a more pronounced EMT phenotype, evident from the significantly higher protein and transcript level of the marker Vimentin (Figure 5 B,C, Supplementary Figure 3A), and lower levels of the MET marker E-cadherin only at transcript level (Figure 5 B,C, Supplementary Figure 3A). Following the conformation of the tumorigenic phenotype of spheroids, we monitored the expression of Wnt players β-catenin, SATB1 and SATB2 in both 2D cell lines and spheroids. The rationale for investigating the expression profile of Wnt players in a tumorigenic system lies in the crucial role played by Wnt/ β-catenin signaling in tumor initiation and progression. It is well established that β-catenin and SATB1 are upregulated in various Wnt driven cancer types including CRC (24). In our results, the transcript levels of β-catenin showed no significant alteration; however, dynamic reciprocal expression in SATB proteins was observed, with SATB1 showing higher levels in spheroids (Figure 5B).

In addition to the transcript levels, SATB1 was also upregulated at the protein level in 3D spheroids compared to 2D cultured cells, while SATB2 levels were downregulated (Figure 5 D). This observation is crucial for elucidating the dynamics between SATB proteins in the context of tumor progression. Given the established fact that SATB1 is upregulated in tumor tissues (22, 59)(Supplementary Figure 5A,B), the increased expression of SATB1 in 3D spheroids mirrors the trend observed in our preliminary study using tissue samples (Supplementary Figure 5C).

Notably, we observed an intriguing correlation between the expression of Vimentin and the upregulation of SATB1 in 3D spheroids, as well as a connection between E-cadherin and SATB2 in 2D cells. This implies a possible association between the EMT-MET markers and the dynamic expression of SATB proteins. Consequently, it can be inferred SATB1 expression may coincide with the EMT phenotype, while SATB2 might be correlated with the MET phenotype. A study in breast cancer stem cells reported a similar correlation, wherein SATB1 led to the upregulation of Snail1 and Twist1 through the activation of Notch signaling (62). Conversely, SATB2 has demonstrated variable effects on the generation and regulation of stem cell or progenitor-like cells in CRC (31,63). Given that the maintenance of stemness and de-differentiation is a hallmark of tumorigenesis, our inferences in context of both SATB1-SATB2 and EMT-MET phenotypes may contribute to an improved understanding of the role of their dynamic expression in epithelial and mesenchymal transitions.

Having established the 3D model system, we performed a colony formation assay to assess the impact of statin treatment on the clonogenicity of CRC cells (Figure 6A). The results showed a significant reduction in the number of colonies upon statin treatment, reaffirming its anti-tumor effect. Collectively, these results demonstrate that 3D spheroids derived from CRC cell lines exhibit distinct expression pattern of epithelial and mesenchymal markers and SATB family proteins.

**Figure 6:**
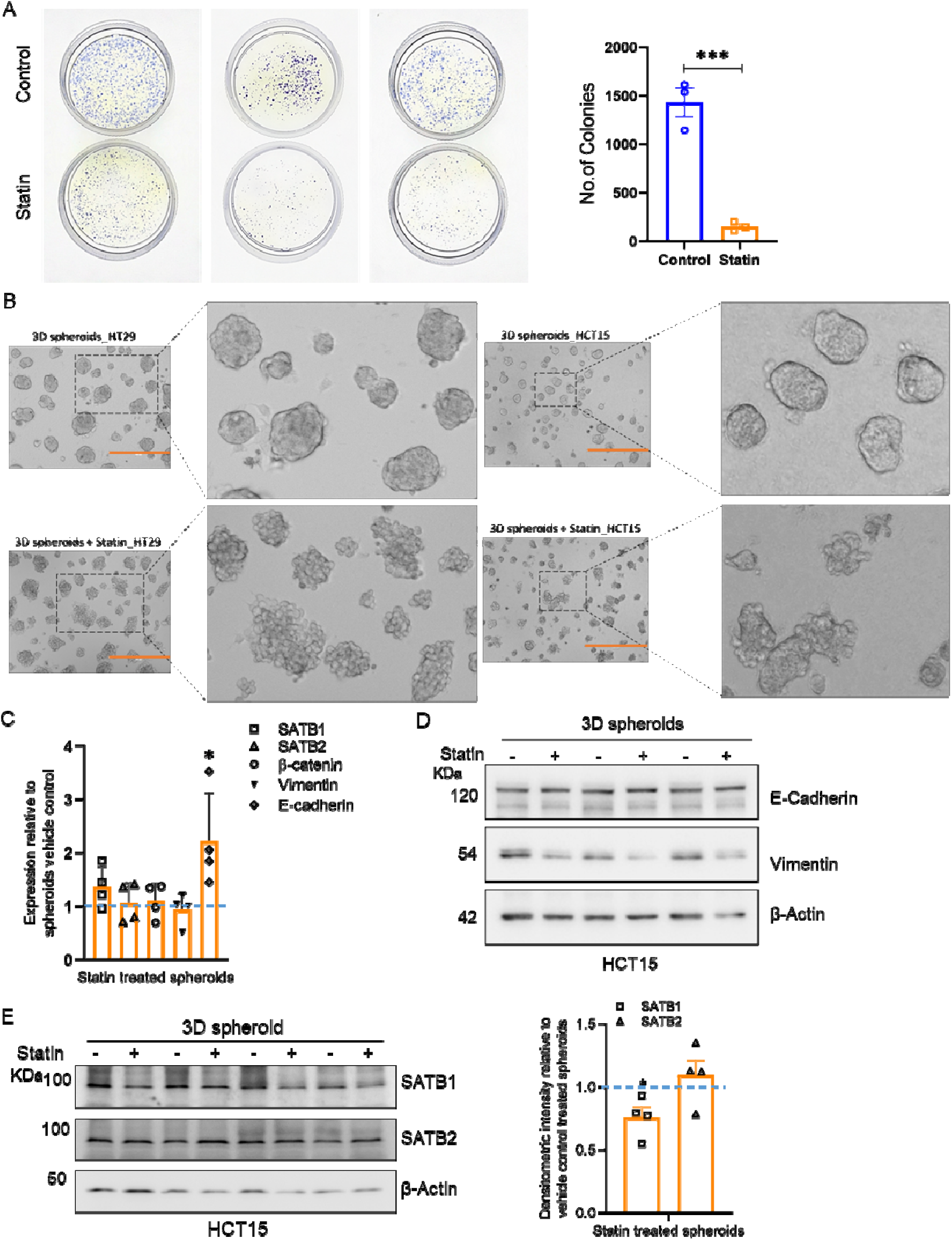
Statin treatment results in reduced SATB1 protein expression and increased E-cadherin expression in spheroids. (A) Colony formation assay of HCT15 cells on statin treatment and graph representing no. of colonies in control versus treated sets. The colonies were allowed to form on soft agar for 5 days and thereafter treated with statin for additional 48 h. The no. of colonies was observed to be significantly reduced on statin treatment. (B) Images to show the effect of statin on HT29 and HCT15 spheroids, where the treated set show disintegration of the spheroids. Phase contrast images, scale bar 400 μm (orange) with zoomed in image in adjacent square box clearly depict a disintegrated spheroid morphology upon statin treatment. The spheroids were allowed to form for 4 days, thereafter statin treatment was given for additional 48 h. (C) Relative expression at transcript levels of SATB1, SATB2, β-catenin in spheroids with and without statin treatment recapitulate the 2D cell line results of no alteration at transcript level. However, the E-cadherin transcript levels are significantly upregulated upon statin treatment, suggesting a transition to an epithelial phenotype. (D) Immunoblot to validate the protein expression of EMT marker Vimentin and MET marker E-cadherin in untreated and statin treated spheroids respectively. The Vimentin protein levels are significantly reduced upon statin treatment. (E) Immunoblot to monitor the protein levels of SATB1 and SATB2 in statin-treated spheroids. The densitometry graphs on right depict the normalized intensities of SATB1 and SATB2. SATB1 levels were significantly reduced, whereas, SATB2 was not significantly altered upon statin treatment. Biological replicates n=4, *p<0.05, ns is non-significant as per Student’s t-test analysis.

### Statin treated 3D spheroids exhibit reversal of the expression patterns of EMT-MET markers and SATB proteins

Next, we treated spheroids derived from the CRC cell lines HT29 and HCT15 with statin, and their morphology was observed. The treated spheroids exhibited disintegration in both the HT29 and HCT15 cell lines in 3D cultures (Figure 6B). Consequently, the effect of statin on both colonies and 3D spheroids prompted further exploration of transcript and protein alterations, particularly regarding the equilibrium in expression levels between the epithelial and mesenchymal markers and the dynamic expression of SATB1 and SATB2 in the spheroid model system. Therefore, we investigated the status of EMT-MET markers in response to statin treatment and found no alterations in Vimentin transcript levels. However, intriguingly the epithelial marker E-cadherin exhibited a significant upregulation upon statin treatment (Figure 6C). However, immunoblots of extracts from 3D spheroids of the HCT15 cell line demonstrated a significant reduction in the protein level of Vimentin upon statin treatment, suggesting a potential shift in phenotype from mesenchymal to epithelial following statin (Figure 6D, Supplementary Figure 3B). Moreover, an inverse effect was observed on the dynamic expression of SATB proteins as statin treatment led to significant downregulation of SATB1, while SATB2 expression remained largely unaltered (Figure 6E). The transcript levels of the both SATB1 and SATB2 remained unaltered (Figure 6C).

In summary, findings from spheroid experiments suggest a potential link between elevated tumorigenicity (representing a more mesenchymal state) and SATB1 expression, as well as reduced tumorigenicity (reflecting a more epithelial state) and SATB2 expression. Nonetheless, treatment with statins seems to trigger a reversal by altering the expression patterns of both SATB proteins and the markers associated with mesenchymal and epithelial phenotypes, which are typically indicative of tumor advancement.

### Statins reduce tumor burden *in vivo* and recapitulate the effect on SATB proteins

Drawing from our spheroid tumorigenic model, we observed pronounced dynamics of expression of SATB1 and SATB2 in conjunction with the EMT-MET phenotype, which was reversed upon statin treatment. Considering that β-catenin did not exhibit significant alterations in expression in both 2D cell lines and their respective 3D spheroids, we proposed that, in our study, β-catenin expression may not contribute to the tumorigenicity as drastically as the dynamic behavior of the SATB proteins. Consequently, we decided to investigate the tumorigenic potential of the reciprocal expression profile of SATB1 and 2 *in vivo*.

Furthermore, we aimed to validate the anti-tumorigenic potential of statins by assessing their impact on the dynamic expression profiles of SATB proteins in both murine and human studies. The murine experimental approach involved subcutaneously injecting CRC cell lines and monitoring tumor burden after 45 days. Statin was orally administered at the dosage of 40 mg/kg/day, initiated 7 days post-cell injection (Figure 7A). Following the statin regimen, tumor weights were recorded and tissues were utilized for transcript and protein analysis.

**Figure 7:**
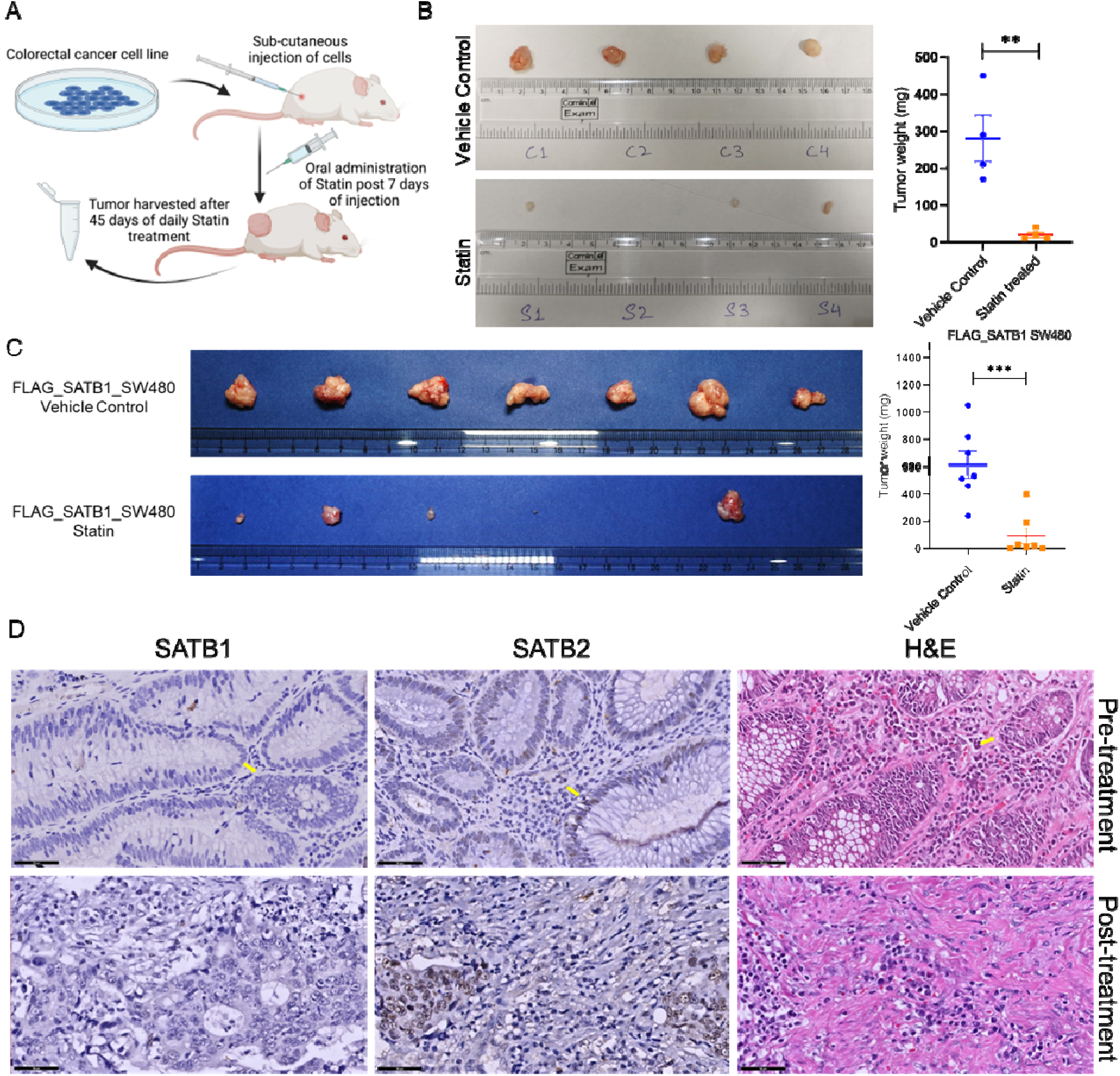
Statin mediated reduction in tumor burden via downregulation of SATB1 *in vivo*. (A) Schematic presentation of the experimental strategy using NOD-SCID mice. (B) Comparative images of the tumors and graph to show a reduction in tumor burden on statin treatment in mice injected with HCT116 cell line, (Biological replicates n=4) (C) FLAG-SATB1 overexpressed SW480 cells were injected in NOD-SCID mice, and tumor burden was observed (n=7) **p<0.005, ***p<0.0005 as per Student’s t-test analysis. (D) The clinical phase II/III trial conducted in rectal patients to incorporate statins in the therapeutic regime. The human study was carried out in collaboration with the TMH, Mumbai gastro-oncology and pathology department. Representative image of molecular profiling of the biopsy samples obtained for immunohistochemistry analysis of SATB1 and SATB2 expression in a paired pre-treatment and post-treatment biopsy section (50 μm scale bar). The yellow arrows in the upper panel, i.e., pre-treatment samples point at the location of the expression signal. For SATB1, the expression is at the adenomatous section, whereas the SATB2 expression is marked at the normal crypt section. The H&E for the sample shows tumorigenic growth. The lower panel i.e., post-treatment section shows mostly stroma with scattered cells as the tumor was scanty and reduced. SATB2 expression seems to be higher in the post-treatment section, whereas, the SATB1 expression is significantly downregulated.

Remarkably, statin-treated mice exhibited a reduced tumor burden compared to the vehicle control in the mice injected with the aggressive CRC cell line HCT116 (Figure 7B). Given our proposal that SATB1 expression correlates with higher tumorigenicity, we sought to validate whether statin treatment in mice aligned with the molecular *in vitro* results. For this purpose, we injected less aggressive CRC cell line, SW480, with a stably incorporated FLAG-tagged overexpression of SATB1 (Supplementary Figure 4A). Following the experimental regime, the tumor burden was significantly reduced upon statin treatment (Figure 7C). Tissues were then isolated and examined for SATB1 and SATB2 expression at both transcript and protein levels. The transcript levels showed no change, consistent with previous findings (Supplementary Figure 4B). However, there appeared to be a slight downregulation of SATB1 in tumor cells and a modest upregulation of SATB2 at the protein level (Supplementary Figure 4C).

To further affirm the potential of the SATB family of proteins as markers for tumor progression and viable targets for statin therapeutics, we designed a human study. We conducted a phase II/III randomized trial approved by the Institutional Ethics Committee (IEC), in collaboration with Tata Memorial Hospital, Mumbai, India, focusing on the inclusion of statin in the treatment of colorectal cancer patients. Patients who were not on prior regimen of statins or aspirin were included in this study. Their rectal adenocarcinoma status was confirmed through initial biopsies. These patients were divided into two treatment arms: Arm ‘A’ received Rosuvastatin in combination with neoadjuvant chemo-radiotherapy, while arm ‘B’ received only the neoadjuvant chemo-radiotherapy. Statins were prescribed at a daily dosage of 20 mg for 12-16 weeks, followed by tumor progression analysis. Chemo-radiotherapy was administered to these patients after completion of statin treatment.

The study encompassed 60 patients, with 30 in arm ‘A’ and 30 in arm ‘B’. Additionally, we examined paired biopsy samples, pre- and post-statin treatment, from 10 out of the 30 patients in arm ‘A’. Patients in arm ‘A’ generally exhibited superior responses to the therapy regimen, displaying partial or near-complete tumor regression. Post-treatment samples indicated sparse and significantly reduced tumors. Out of the 30 patients in Arm A, 48% displayed tumor regression and showed no signs of relapse during the one-year follow-up. Conversely, 28% of the patients not receiving statins in arm B exhibited a response to therapy, with higher incidences of relapse.

Molecular study of SATB1 and SATB2 expression profiles was conducted by observing through Immunohistochemistry (IHC) analysis on tissue samples from the paired biopsies of these 10 patients. The IHC analysis revealed a marked reduction in SATB1 staining in post-treatment samples compared to their paired pre-treatment counterparts (Supplementary Table 5). SATB2 expression, however, showed variability between the two conditions. A representative image of one patient’s pre- and post-treatment SATB1/2 expression profiles (Figure 7D, Supplementary Figure 6) demonstrated diminished SATB1 expression on statin treatment.

The distinct expression profiles of SATB1 and SATB2 proteins in the rectal samples further reinforce our proposition for their clinical recognition as tumorigenic markers and targets for the repurposed drug statins. Understanding the specific mechanism by which statins regulate these key players in the Wnt pathway holds significance for colorectal cancer therapeutics.

## Discussion

Since its discovery in the 1970s, the Statin family of drugs has represented a groundbreaking advancement in the treatment of hyperlipidemia (14). Serving as competitive inhibitors of the rate-limiting step in the mevalonate pathway, these drugs have revolutionized the approach to therapeutics for conditions such as atherosclerosis, fatty liver, and coronary plaques (15,17). The unique lactone ring structure of statins acts as a substrate for HMG-CoA reductase, preventing the production of mevalonate and consequently inhibiting downstream products, including cholesterol and its derivatives. Statins can be categorized into two types based on the R group attached to the lactone ring: hydrophobic and hydrophilic (64). The R groups play a crucial role in drug delivery and statin’s processivity within the cell. Hydrophobic statins like simvastatin, fluvastatin, pitavastatin, lovastatin, and atorvastatin are known for their easy cell penetration, while hydrophilic statins like rosuvastatin and pravastatin become more effective after metabolism in the liver (64). The diverse R groups, providing hydrophobicity and hydrophilicity, also influence the pleiotropic side effects observed in early formulations (16). The first commercially available statin, lovastatin, was associated with side effects such as nausea, fatigue and constipation, which were progressively reduced in subsequent statins like simvastatin and pravastatin (65). The synthetically produced statins like rosuvastatin and atorvastatin effectively addressed these side effects (65). In our study we employed both simvastatin and rosuvastatin, achieving similar effects in both *in cellulo* and *in vivo* systems. However, the human study specifically utilized rosuvastatin.

Over the past two decades, the consideration of potential anti-neoplastic properties of statins has been fueled by their effect on the cholesterol pathway, which deprives tumor cells of essential lipids (66-68). While previous studies have emphasized the apoptotic and cell cycle targeting effects of statins on tumorigenesis (69-71), another report suggested specificity of statin effect on APC mutated CRC cells as well as APC mutant patient derived xenografts (72). Our investigation focused on the statin-mediated influence on pathways responsible for early adenoma formation in colorectal cancer (CRC). Published reports have highlighted a strong correlation between elevated cholesterol levels and the upregulation of Wnt signaling (73). In our transcriptome and proteome data, we observed a specific targeting of Wnt-responsive genes with statin treatment. Mechanistically, two major players of the Wnt pathway, β-catenin and SATB1, were downregulated at the protein level rather than the transcript level. This suggests that statins may impact the stability of these proteins, hinting at a degradation process.

Furthermore, we proposed that the statin effect on Wnt signaling might rely on mevalonate pathway inhibition, as evidenced by the rescue in the protein levels of β-catenin and SATB1 upon mevalonate supplementation. Additionally, our proteome data revealed a decrease in YAP expression, corroborating with recent findings reporting YAP downregulation in response to statin treatment, dependent on the cholesterol pathway inhibition (74-76). Hence, our comprehensive analysis of statin mediated effect on the early stages of tumorigenesis not only validated established molecular targets but also revealed novel targets of statin treatment, such as the SATB family of proteins, recognized as major players in Wnt/ β-catenin signaling.

The effect of statin treatment on chromatin organizer SATB proteins is pivotal, given their role in regulating the expression profile of downstream Wnt target genes. Reports indicate that SATB1 specifically contributes to the upregulation of Wnt responsive genes in conjunction along with β-catenin and TCF/LEF transcription factors (22). The reduction in SATB1 protein levels due to statin treatment directly implies diminished tumorigenicity. Conversely, SATB2, known for its reciprocal expression pattern, acts as a tumor suppressor (26). Although statin treatment did not exhibit a significant effect on the SATB2 levels as pronounced as SATB1, the dynamic expression of both SATB proteins was disrupted in tumorigenic tissue and 3D spheroids. Moreover, survival analysis of patients with various cancer types indicated that the expression of SATB family chromatin organizers, specifically SATB1 and SATB2, is correlated with tissue-specific expression patterns and may influence disease prognosis (24).

3D spheroids emerged as a superior model system compared to 2D cell lines for studying the reciprocal expression of SATB1 and SATB2, validated by EMT-MET markers indicative of increased tumorigenic potential. Strong correlations were observed between higher SATB1 and Vimentin levels in spheroids, and elevated SATB2 and E-cadherin expression in the corresponding 2D cells. Statin treatment in spheroids not only reversed the SATB protein expression but also led to a higher transcript level of E-cadherin, implying a shift toward an epithelial phenotype. Morphologically, the spheroids underwent disintegration upon statin treatment, indicating the initiation of a phenotype switch. Taken together, the interplay between SATB protein expression, EMT-MET phenotype, and the tumorigenic versus anti-tumorigenic activity of statins suggests a regulatory mechanism. Exploring the relationship between SATB chromatin organizers and EMT-MET transitions, both with and without statin treatment, becomes intriguing, with a focus on the potential role of SATB proteins as EMT-MET markers encompassing tumor progression (24).

The *in vivo* analysis performed using mice and the examination of biopsy samples obtained from CRC patients further validated the statin mediated effect on SATB proteins. NOD-SCID mice, subcutaneously injected with HCT116 cells and the SW480 FLAG-SATB1 overexpressing cells, respectively, exhibited a significant reduction in tumor burden on statin treatment, accompanied by decreased SATB1 expression, mirroring *in cellulo* observations. In the ongoing human study, a randomized phase II/III trial in collaboration with Tata Memorial Hospital, Mumbai, our preliminary findings from 60 equally divided patients in Arms A (30 patients receiving statin treatment with neoadjuvant therapy) and B (30 patients receiving only neoadjuvant therapy) indicate a better response in Arm A, emphasizing on improved outcomes with the addition of statins to the regimen.

Furthermore, molecular profiling of SATB protein expression was performed using the histopathological samples obtained from 10 patients in arm ‘A’. Analysis of paired biopsies from pre-treatment versus post-statin treatment regime revealed reduced SATB1 expression in tumor sections. SATB2 expression varied in immunohistochemistry analysis, but the signal intensity distribution appeared higher in normal tissue than in tumorigenic adenomatous tissue, as confirmed morphologically by H&E staining (Figure 7D, yellow arrow). The results may provide valuable information regarding the potential benefits of the incorporating statins into the current therapeutic regimen, leveraging their anti-neoplastic mechanism targeting SATB proteins.

Despite numerous reported players, characterizing molecular markers for aggressive diseases remains a significant challenge. However, SATB proteins exhibit consistent expression profiles in both *in vitro* and *in vivo* settings. Additionally, the substantial impact of statins on the Wnt pathway via SATB proteins underscores the significance of our results. Therefore, our findings add to the expanding repertoire of molecular targets for prognosis and treatment, enhancing our understanding of the repurposed function of statins in colorectal cancer therapeutics.

## Materials and Methods

### Cell culture and spheroid formation

HCT116 and HCT15 CRC cell lines were obtained from the European Collection of Cell Cultures (ECACC) and HT29 cell line was obtained from American Type Culture Collection (ATCC, Manassas, Virginia, USA). HCT116 was cultured in DMEM media (Gibco, Waltham, MA, USA) with 10% FBS whereas HCT15 and HT29 were cultured in RPMI media (Gibco, Grand Island, NY, USA) with 10% FBS. Spheroid formation was performed according to the protocol, as described (77). Briefly, the 24-well dish was coated with matrigel bed and allowed to solidify for 30 min at 37°C. Single-cell suspension of the cells was made and 10^4^ cells were seeded per well. On-top matrigel was added in media and spheroids were allowed to form for 4 days. Colony formation assay was also performed as described (78). Briefly, 1.2% low melting point agarose was used as the base with media. 10^4^ cells were seeded in a 60 mm dish and topped with 0.6% agarose + media. The colonies were allowed to grow for 7 days in total. Matrigel was obtained from Corning (Cat. No. 356231). Simvastatin and Rosuvastatin used in all assays were obtained from Sigma (Simvastatin Cat No. 79902-63-9, Rosuvastatin #SML-1264). Rosuvastatin used for *in vivo* assays was obtained from Cipla (Rosulip).

### Western blotting and RNA isolation

The antibodies used were as follows, β-catenin (BD Bioscience Cat No. 610153), SATB1_L745 (Cell Signaling Tech, Cat No. 3650), SATB2 (AbCam, Cat No. Ab92447), GAPDH (AbCam, Cat No. G041), TCF4 (CST #2569S), AXIN2 (CST #76G6), PCAF (Santa Cruz #13124), β-Actin (Sigma #A2228), E-cadherin (AbCam #ab40772), Vimentin (AbCam #ab92547). The lysates were prepared in RIPA buffer (20 mM Tris pH7.5, 150 mM NaCl, 1 mM EDTA, 1 mM EGTA, 1% NP-40, 1% Sodium deoxycholate, protease inhibitor cocktail (PIC, Thermo Fisher Cat No. A32963). Total RNA was extracted from cells grown under 2D and 3D culture conditions and vehicle control and statin-treated sets using Trizol reagent (Taraka, Cat No. 9109). Extracted RNA was either subjected to library preparation for high-throughput sequencing or PCR-based gene expression analysis. For qPCR-based gene expression analysis, cDNA was synthesized using High-capacity cDNA Reverse Transcription kit (Applied Biosystems Cat No. 4368814). The cDNA was utilized for relative gene expression analysis using gene-specific primers, Supplementary table, with 18srRNA as endogenous control.

### Transcriptome analysis sample preparation

Total RNA (500 ng) was subjected to mRNA purification using NEBNext Poly(A) mRNA Magnetic Isolation Module (NEB, US) according to the manufacturer’s instructions. The purified mRNA was used for library preparation using NEBNext Ultra II RNA Library Prep Kit for Illumina (NEB, US) using the protocol provided in the kit. The final libraries were purified using HighPrep PCR Clean-up System (MagBio Genomics, USA) and were quantified using the Qubit 1X HS DNA system (Thermo Fisher Scientific). All the libraries were pooled in equimolar ratios and subjected to 75 bp PE chemistry on Nextseq550, Illumina.

### RNAseq analysis

Paired-end sequencing was performed using RNA samples on Illumina platform (Macrogen Inc, Korea). In brief, after performing quality control (QC), qualified samples were processed for library construction. Sequencing library was prepared by random fragmentation of cDNA followed by adapter ligation. Adapter ligated fragments were PCR amplified and gel purified. The libraries were loaded into a flow cell and each fragment was clonally amplified through bridge amplification. The sequencing data was converted into raw data for analysis. The files have been submitted to SRA database and the accession number for the same is PRJNA957223.

The quality of sequencing reads was checked using FASTQC (version 0.10.1) [ Andrews, S. (2010). FastQC: a quality control tool for high throughput sequence data] Available online at: http://www.bioinformatics.babraham.ac.uk/projects/fastqc/]. The sequence alignment was performed on the human genome (version hg38) [https://www.gencodegenes.org/] using HiSAT2 (v 2.05) [https://daehwankimlab.github.io/hisat2/manual/]. The resulting bam files were used to generate a count matrix (85) followed by differential expression analysis (v 1.18.1) (86). Ontology analysis was carried out using clusterProfiler (v 3.6.0) (87,88).

### Lipidomics analysis

All the samples for lipidomics analysis were prepared and analyzed using established protocols previously described by us (79-81) on a Sciex X500R QTOF mass spectrometer, fitted with an Exion series UHPLC. Briefly, statin treatment was given in the CRC cell line HCT116 and cells were harvested for lipid extraction. Chloroform and methanol (2:1) were used to process the samples. The lipids were run in negative and positive modes in the LC-MS method, keeping Free Fatty Acid (FFA 17:1, 1 nanomole final concentration) as the internal standard.

### Proteomics analysis

Whole cell lysates were resolved on a 10% SDS-PAGE gel and subsequently processed for proteomics using standard in-gel sample preparation protocols (82). Briefly, the gel bands of interest were excised, destained, reductively alkylated and digested using MS-grade trypsin. Subsequently, the peptides were desalted and processed for proteomics analysis on Sciex TripleTOF6600 mass spectrometer interfaced with an Eksigent nano-LC 425 system using established protocols previously reported by us (83,84). All raw data was analyzed using the ProteinPilot software from Sciex using search parameters previously described by us (84). The GO term for the differential peptides were plotted using g:Profiler. The mass spectrometry proteomics data have been deposited to the ProteomeXchange Consortium via the PRIDE [1] partner repository with the dataset identifier PXD041210.

Reviewer account details:

Username

Password: 0dl2bhOo

### *In vivo* study of tumor in mice

HCT116 cells and SW480-FLAG overexpressing SATB1 cells, SW480-FLAG control cells (1 × 10^6^ each) were separately injected subcutaneously in 8 NOD-SCID mice (male or female respectively, 6-8 weeks old, weighing 25-30 g). They were equally divided into two sets, where one set was designated vehicle control and the other set was given statin treatment. Simvastatin (given to mice injected with HCT116 cells) /Rosuvastatin (given to mice injected with SW480 SATB1 overexpressing cells) were dissolved in 0.5% methyl-cellulose and administered orally on a daily basis for 45 days, post-7 days of injection of cells. The dose provided was 40 mg/kg/day and no side effects were observed. The mice were sacrificed after 45 days of the assay and the tumor was isolated and weighed. All the murine experiments were approved by the Institutional Animal Ethics Committee.

### Clinical trial (IECHR/VB/2018/014)

The phase II clinical trial was conducted at Tata Memorial Hospital (TMH), Parel, Mumbai, India under ethical approval by the Institutional Ethics Committee for Human Research, after a written informed consent from the patients. The paired samples of the rectal tumor and adjacent normal tissues were obtained from the rectal cancer patients participating in the study. The major aim of the study is to validate whether statins show a better pathological complete response (pCR) in combination with Neo-adjuvant chemo-radiotherapy (NACTRT). The secondary aims include the overall comparison and survival of the patients over a period of 5 years, in the two arms of the study, one with statin added to NACTRT (arm A) and the other with only NACTRT (arm B). Rosuvastatin (20 mg/day) was provided to the patients in arm A, with NACTRT for 6 weeks, followed by only Rosuvastatin (20 mg/day) for another 6-10 weeks, until the surgery, summing up to a total of 12-14 weeks of rosuvastatin treatment. The patients were followed up for approximately 1 year for relapses and survival.

The inclusion criteria for the randomized trial for receiving rosuvastatin were carefully structured. The patients should be between the age group of 18 to 70, and should be willing to consent to the study. They should be confirmed for adenocarcinoma of the rectum and should be fit for receiving neoadjuvant chemotherapy. The disease should not be metastatic and there should be no history of Crohn’s disease or ulcerative colitis. Patients should not be on any medication reactive to statin or prior aspirin consumption. Female patients should not be nursing or pregnant. All the patients were recorded for any comorbidities.

We have recorded 60 patients so far, abiding by all the criteria. The percentage of patients showing better prognosis in the follow-up routines were manually calculated for these 60 patients. 30 patients belonged to arm A with statin in their regime and 30 in arm B with only the NACTRT. We could also conduct immunohistological study in 10 patients from arm A, with paired biopsy samples from pre-statin treated condition and post-statin treated condition.

The tissue sample collection was procured with standard endoscopic and surgical procedures. Immunohistochemistry was performed for the tissue biopsies embedded in paraffin blocks and probed using SATB1 (Santa Cruz #sc-376096) and SATB2 (Abcam #ab92446 anti-SATB2) antibodies. H score was calculated with intensity levels, weak = 1, moderate = 2, strong = 3 and % of cells positive for signal, 0-33% = 1, 34-66% = 2 and 64-100% = 3. The score for intensity was multiplied with score for percentage of cells to give the final H score. Tissue samples for protein expression assays were collected and snap-frozen for shipment in dry-ice. The tissues were immediately homogenized in RIPA buffer and lysates were run for 11 pair of normal adjacent and tumor samples (Supplementary Figure 5C).

### Statistical Analysis

All the experiments were performed in biological triplicates. The statistical analysis between two groups were performed using the two-tailed Student t-test unpaired method. Unless, otherwise mentioned, 0.05 level of confidence was accepted for statistical significance.

## Declarations

### Funding

ST acknowledges support from IISER Pune, Scivic Engineering India Pvt Ltd and the Innoplexus Consulting Services Pvt. Ltd., and Department of Biotechnology (DBT) for fellowships. EG acknowledges support from the Prime Minister’s Research Fellowship. The authors thank Ankita Sharma for help with RNA-seq library preparation. The clinical trial (Project no 3033 received intramural funding from the Tata Memorial Center). The work was supported by research grant from Department of Biotechnology (DBT), Government of India to SG, PP, and AS (BT/ATGC/127/SP39484/2020). SG is also a recipient of the JC Bose Fellowship (JCB/2019/000013) by the Science and Engineering Research Board, Government of India. Authors wish to thank Department of Science and Technology (DST)-FIST for supporting the establishment of the mass spectrometry facility (grant number SR/FST/LSII-043/2016) in the Department of Biology, IISER Pune and a SwarnaJayanti fellowship of SK (grant number SB/SJF/2021-22/01). We thank the National Facility for Gene Function in Health and Disease (supported by a grant from the Department of Biotechnology, Government of India; BT/INF/<u>22/SP17358/2016</u>) at IISER Pune for maintaining and providing mice for this study.

### Ethical approval

The research on animal models was conducted in accordance with the regulations of the Institutional Animal Ethics Committee and was approved by the same. Research on human subjects was approved by the Institutional Ethics Committee for Human Research (IISER Pune) and the human trial study was approved by the Institutional Review Board (Tata Memorial Hospital) in accordance with the Declaration of Helsinki. All patients were enrolled in the study after written informed consent.

### Competing Interests

The authors declare no conflict of interest.

### Authors’ contributions

The project was conceived by SG, RN, and ST. The experimental design was contributed by SG, RN, ST, RM, Satyajeet K, and SK. The experiments were performed by ST and EG. The clinical trial was conceived by PP, AS and SG. The biopsies were provided from patients enrolled on the study with the help of SD. IHC was performed by SH, SY, MB. The manuscript was written by ST and SG.

### Availability of data and materials

The mass spectrometry proteomics data have been deposited and can be accessed at the ProteomeXchange Consortium via the PRIDE [1] partner repository with the dataset identifier PXD041210.

Reviewer account details:

Username

Password: 0dl2bhOo

### Supplementary Data statement

Supplementary Data are available at NAR Online.

## Acknowledgments

The authors acknowledge the funding agencies for supporting the work. We thank Ankita Sharma for help with RNA-seq library preparation. We thank the National Facility for Gene Function in Health and Disease (supported by a grant from the Department of Biotechnology, Government of India; BT/INF/<u>22/SP17358/2016</u>) at IISER Pune for maintaining and providing mice for this study. We also acknowledge Dr. Jasmine Kaur Dhall for help with the processing of human tissue biopsies and immunoblot analysis.

## Notes

### Competing Interest Statement

The authors have declared no competing interest.

