## Supplementary Information for "Statins attenuate Wnt/β-catenin signaling by targeting SATB family proteins in colorectal cancer"

### Supplementary Data:

| ID | Source | Term ID | Term Name | $p_{adj}$ (query_1) |
|---|---|---|---|---|
| 1 | GO:BP | GO:0001676 | long-chain fatty acid metabolic process | $2.214\times10^{-3}$ |
| 2 | GO:BP | GO:0006082 | organic acid metabolic process | $1.894\times10^{-4}$ |
| 3 | GO:BP | GO:0006631 | fatty acid metabolic process | $1.147\times10^{-4}$ |
| 4 | GO:BP | GO:0008299 | isoprenoid biosynthetic process | $5.599\times10^{-5}$ |
| 5 | GO:BP | GO:0010033 | response to organic substance | $1.691\times10^{-5}$ |
| 6 | GO:BP | GO:0019216 | regulation of lipid metabolic process | $9.291\times10^{-8}$ |
| 7 | GO:BP | GO:0006629 | lipid metabolic process | $1.334\times10^{-8}$ |
| 8 | GO:BP | GO:0008610 | lipid biosynthetic process | $1.505\times10^{-9}$ |
| 9 | GO:BP | GO:0019218 | regulation of steroid metabolic process | $1.208\times10^{-9}$ |
| 10 | GO:BP | GO:0006066 | alcohol metabolic process | $3.867\times10^{-13}$ |
| 11 | GO:BP | GO:0006694 | steroid biosynthetic process | $6.240\times10^{-14}$ |
| 12 | GO:BP | GO:0008202 | steroid metabolic process | $1.721\times10^{-15}$ |
| 13 | GO:BP | GO:0008203 | cholesterol metabolic process | $2.319\times10^{-17}$ |
| 14 | GO:BP | GO:0006695 | cholesterol biosynthetic process | $1.378\times10^{-19}$ |
| 15 | GO:BP | GO:0016125 | sterol metabolic process | $1.678\times10^{-19}$ |
| 16 | GO:BP | GO:0045540 | regulation of cholesterol biosynthetic process | $4.569\times10^{-13}$ |
| 17 | GO:BP | GO:0046165 | alcohol biosynthetic process | $1.549\times10^{-12}$ |
| 18 | GO:BP | GO:0090181 | regulation of cholesterol metabolic process | $6.932\times10^{-12}$ |
| 19 | GO:BP | GO:0106118 | regulation of sterol biosynthetic process | $4.569\times10^{-13}$ |
| 20 | GO:BP | GO:1902652 | secondary alcohol metabolic process | $4.145\times10^{-17}$ |
| 21 | GO:BP | GO:1901617 | organic hydroxy compound biosynthetic process | $2.775\times10^{-13}$ |
| 22 | GO:BP | GO:0046890 | regulation of lipid biosynthetic process | $7.293\times10^{-10}$ |
| 23 | GO:BP | GO:0050810 | regulation of steroid biosynthetic process | $9.586\times10^{-11}$ |
| 24 | GO:BP | GO:0044283 | small molecule biosynthetic process | $1.603\times10^{-9}$ |
| 25 | GO:BP | GO:0044281 | small molecule metabolic process | $2.021\times10^{-8}$ |
| 26 | GO:BP | GO:0044255 | cellular lipid metabolic process | $1.758\times10^{-5}$ |
| 27 | GO:BP | GO:0033993 | response to lipid | $4.828\times10^{-4}$ |
| 28 | GO:BP | GO:0019752 | carboxylic acid metabolic process | $3.530\times10^{-5}$ |
| 29 | GO:BP | GO:0046949 | fatty-acyl-CoA biosynthetic process | $1.644\times10^{-4}$ |
| 30 | GO:BP | GO:0071616 | acyl-CoA biosynthetic process | $5.229\times10^{-4}$ |
| 31 | GO:BP | GO:0006637 | acyl-CoA metabolic process | $6.871\times10^{-3}$ |
| 32 | GO:BP | GO:1901568 | fatty acid derivative metabolic process | $1.351\times10^{-5}$ |
| 33 | KEGG | KEGG:01212 | Fatty acid metabolism | $1.176\times10^{-2}$ |
| 34 | KEGG | KEGG:01100 | Metabolic pathways | $1.081\times10^{-5}$ |
| 35 | REAC | REAC:R-HSA-89... | Fatty acid metabolism | $3.210\times10^{-3}$ |
| 36 | REAC | REAC:R-HSA-68... | Cholesterol biosynthesis via lathosterol | $8.372\times10^{-4}$ |
| 37 | REAC | REAC:R-HSA-75... | Fatty acyl-CoA biosynthesis | $2.887\times10^{-4}$ |
| 38 | REAC | REAC:R-HSA-14... | Metabolism | $2.976\times10^{-4}$ |
| 39 | WP | WP:WP4804 | Cholesterol Biosynthesis with Skeletal Dysplasias | $7.289\times10^{-6}$ |
| 40 | WP | WP:WP3963 | Mevalonate pathway | $7.289\times10^{-6}$ |
| 41 | WP | WP:WP1982 | Sterol Regulatory Element-Binding Proteins (SREB... | $6.915\times10^{-7}$ |
| 42 | REAC | REAC:R-HSA-55... | Metabolism of lipids | $4.046\times10^{-9}$ |
| 43 | REAC | REAC:R-HSA-16... | Regulation of cholesterol biosynthesis by SREBP (S... | $1.142\times10^{-10}$ |
| 44 | REAC | REAC:R-HSA-89... | Metabolism of steroids | $7.705\times10^{-12}$ |
| 45 | REAC | REAC:R-HSA-24... | Activation of gene expression by SREBF (SREBP) | $6.627\times10^{-12}$ |
| 46 | WP | WP:WP197 | Cholesterol Biosynthesis Pathway | $1.815\times10^{-16}$ |
| 47 | WP | WP:WP4718 | Cholesterol metabolism (includes both Bloch and ... | $8.528\times10^{-22}$ |
| 48 | REAC | REAC:R-HSA-19... | Cholesterol biosynthesis | $7.511\times10^{-21}$ |
| 49 | KEGG | KEGG:00100 | Steroid biosynthesis | $1.343\times10^{-13}$ |
| 50 | WP | WP:WP2011 | SREBF and miR33 in cholesterol and lipid homeost... | $3.801\times10^{-4}$ |
| 51 | WP | WP:WP357 | Fatty Acid Biosynthesis | $4.500\times10^{-3}$ |
| 52 | WP | WP:WP4724 | Omega-9 FA synthesis | $2.399\times10^{-3}$ |
| 53 | REAC | REAC:R-HSA-45... | Interleukin-2 family signaling | $1.245\times10^{-2}$ |
| 54 | MIRNA | MIRNA:hsa-miR... | hsa-miR-335-5p | $9.831\times10^{-14}$ |
| 55 | TF | TF:M10108 | Factor: WT1; motif: RGGNGGGGGAGGRGGNGGRG | $1.687\times10^{-2}$ |
| 56 | TF | TF:M12160 | Factor: KLF15; motif: RCCMCRCCCMCN | $4.480\times10^{-2}$ |
| 57 | TF | TF:M07289 | Factor: GKLF; motif: NNNRGGNGNGGSN | $4.670\times10^{-4}$ |
| 58 | TF | TF:M07040 | Factor: GKLF; motif: NNRRGRRNGNSNNN | $2.406\times10^{-4}$ |
| 59 | TF | TF:M00915_0 | Factor: AP-2; motif: SNNNCCNCAGGCN; match cla... | $3.099\times10^{-3}$ |
| 60 | TF | TF:M00444 | Factor: VDR; motif: GGGKNARNRRGGWSA | $9.565\times10^{-3}$ |

| | |
|---|---|
| **version** | e100_eg47_p14_7733820 |
| **date** | 8/27/2020, 1:30:12 PM |
| **organism** | hsapiens |

g:Profiler

**Supplementary Table 1**: **GO terms list for the Manhattan plot (Figure 2A) provided from g:Profiler software for the significantly upregulated genes on statin treatment in transcriptome analysis**. The color coding is dependent on the significance value (p value) assigned to the terms in the software based on the gene list belonging to the term and the statistical significance of their match, blue signifying highly significant to orange signifying least significant. Therefore, all the GO terms listed are highly significant.

| ID | Source | Term ID | Term Name | $p_{adj}$ (query_1) |
|---|---|---|---|---|
| 1 | GO:MF | GO:0003677 | DNA binding | $3.816\times10^{-2}$ |
| 2 | GO:MF | GO:0003824 | catalytic activity | $1.268\times10^{-2}$ |
| 3 | GO:MF | GO:0000217 | DNA secondary structure binding | $3.312\times10^{-3}$ |
| 4 | GO:MF | GO:0003676 | nucleic acid binding | $2.715\times10^{-6}$ |
| 5 | GO:MF | GO:0033170 | protein-DNA loading ATPase activity | $1.165\times10^{-3}$ |
| 6 | GO:MF | GO:0003697 | single-stranded DNA binding | $4.434\times10^{-8}$ |
| 7 | GO:BP | GO:0007049 | cell cycle | $1.115\times10^{-22}$ |
| 8 | GO:BP | GO:0022402 | cell cycle process | $2.890\times10^{-20}$ |
| 9 | REAC | REAC:R-HSA-16... | Cell Cycle | $2.733\times10^{-23}$ |
| 10 | REAC | REAC:R-HSA-69... | Cell Cycle, Mitotic | $3.724\times10^{-18}$ |
| 11 | WP | WP:WP2363 | Gastric Cancer Network 2 | $6.292\times10^{-4}$ |
| 12 | WP | WP:WP2446 | Retinoblastoma Gene in Cancer | $2.425\times10^{-13}$ |
| 13 | WP | WP:WP179 | Cell Cycle | $3.092\times10^{-9}$ |
| 14 | TF | TF:M04826_0 | Factor: p300; motif: ACNTCCG; match class: 0 | $2.934\times10^{-2}$ |
| 15 | TF | TF:M03867_0 | Factor: c-Myc; motif: CACGTGGC; match class: 0 | $2.126\times10^{-2}$ |
| 16 | TF | TF:M01145 | Factor: c-Myc; motif: RACCACGTGCTC | $5.714\times10^{-5}$ |
| 17 | TF | TF:M07601_1 | Factor: C-Myc; motif: NGCCACGTGNN; match class... | $2.719\times10^{-5}$ |
| 18 | TF | TF:M01154 | Factor: c-Myc; motif: KACCACGTGSYY | $9.438\times10^{-7}$ |
| 19 | TF | TF:M01145_1 | Factor: c-Myc; motif: RACCACGTGCTC; match class:... | $3.938\times10^{-7}$ |
| 20 | TF | TF:M07206_1 | Factor: E2F-1; motif: NGGGCGGGARV; match class: 1 | $2.744\times10^{-8}$ |
| 21 | TF | TF:M07601_0 | Factor: C-Myc; motif: NGCCACGTGNN; match class... | $5.262\times10^{-9}$ |
| 22 | TF | TF:M11531_1 | Factor: E2F-2; motif: GCGCGCGCGYW; match class: 1 | $5.023\times10^{-10}$ |
| 23 | TF | TF:M04743 | Factor: c-Myc; motif: NSCACGTGGN | $1.897\times10^{-11}$ |
| 24 | TF | TF:M00797_0 | Factor: HIF1; motif: GNNKACGTGCGGNN; match cl... | $3.022\times10^{-4}$ |
| 25 | TF | TF:M07384_0 | Factor: HIF-1alpha; motif: NCACGTNN; match class... | $1.318\times10^{-2}$ |
| 26 | TF | TF:M00322_1 | Factor: c-Myc:Max; motif: GCCAYGYGSN; match cla... | $6.492\times10^{-3}$ |
| 27 | TF | TF:M09992_1 | Factor: c-Myc; motif: NCCACGTGCNN; match class: 1 | $1.086\times10^{-3}$ |
| 28 | TF | TF:M11081_1 | Factor: SREBP-1; motif: RTCRCGTGAY; match class: 1 | $2.337\times10^{-8}$ |
| 29 | TF | TF:M11081 | Factor: SREBP-1; motif: RTCRCGTGAY | $1.624\times10^{-5}$ |
| 30 | TF | TF:M11081_0 | Factor: SREBP-1; motif: RTCRCGTGAY; match class: 0 | $1.624\times10^{-5}$ |
| 31 | TF | TF:M11601 | Factor: TCF-1; motif: ACATCGRGRCGCTGW | $3.351\times10^{-4}$ |
| 32 | TF | TF:M11603_1 | Factor: TCF-1; motif: ACATCGRGRCGCTGW; match ... | $1.993\times10^{-2}$ |
| 33 | TF | TF:M11603 | Factor: TCF-1; motif: ACATCGRGRCGCTGW | $2.831\times10^{-2}$ |
| 34 | MIRNA | MIRNA:hsa-miR... | hsa-miR-215-5p | $1.790\times10^{-11}$ |
| 35 | MIRNA | MIRNA:hsa-miR... | hsa-miR-192-5p | $5.363\times10^{-12}$ |
| 36 | MIRNA | MIRNA:hsa-miR... | hsa-miR-193b-3p | $2.609\times10^{-15}$ |
| 37 | GO:CC | GO:0005657 | replication fork | $6.730\times10^{-9}$ |

| | | |
|---|---|---|
| **version** | e100_eg47_p14_7733820 | g:Profiler |
| **date** | 8/27/2020, 12:54:33 PM | |
| **organism** | hsapiens | |

**Supplementary Table 2**: **GO terms list for the Manhattan plot (Figure 2B) provided from g:Profiler software for the significantly downregulated genes on statin treatment in transcriptome analysis**. The color coding is dependent on the significance value (p value) assigned to the terms in the software based on the gene list belonging to the term and the statistical significance of their match, blue signifying highly significant to orange signifying least significant. Therefore, all the GO terms listed are highly significant.

| ID | Source | Term ID | Term Name | $p_{adj}$ (query_1) |
|---|---|---|---|---|
| 1 | GO:MF | GO:0045296 | cadherin binding | $9.638\times10^{-12}$ |
| 2 | GO:MF | GO:0050839 | cell adhesion molecule binding | $1.710\times10^{-8}$ |
| 3 | GO:MF | GO:0097159 | organic cyclic compound binding | $2.638\times10^{-4}$ |
| 4 | GO:MF | GO:1901363 | heterocyclic compound binding | $3.704\times10^{-4}$ |
| 5 | GO:MF | GO:0031625 | ubiquitin protein ligase binding | $2.489\times10^{-2}$ |
| 6 | GO:BP | GO:1901575 | organic substance catabolic process | $1.332\times10^{-10}$ |
| 7 | GO:BP | GO:0043161 | proteasome-mediated ubiquitin-dependent protei... | $1.756\times10^{-5}$ |
| 8 | GO:BP | GO:0010498 | proteasomal protein catabolic process | $1.065\times10^{-4}$ |
| 9 | GO:BP | GO:0051603 | proteolysis involved in cellular protein catabolic pr... | $4.716\times10^{-4}$ |
| 10 | GO:BP | GO:0006511 | ubiquitin-dependent protein catabolic process | $5.869\times10^{-3}$ |
| 11 | GO:BP | GO:0017015 | regulation of transforming growth factor beta rece... | $6.255\times10^{-3}$ |
| 12 | GO:BP | GO:1903844 | regulation of cellular response to transforming gr... | $7.426\times10^{-3}$ |
| 13 | GO:BP | GO:0070498 | interleukin-1-mediated signaling pathway | $1.620\times10^{-2}$ |
| 14 | GO:BP | GO:0016055 | Wnt signaling pathway | $3.712\times10^{-2}$ |
| 15 | GO:BP | GO:0198738 | cell-cell signaling by wnt | $3.870\times10^{-2}$ |
| 16 | REAC | REAC:R-HSA-21... | Downregulation of TGF-beta receptor signaling | $6.832\times10^{-7}$ |
| 17 | REAC | REAC:R-HSA-89... | Regulation of PTEN localization | $3.356\times10^{-6}$ |
| 18 | REAC | REAC:R-HSA-21... | TGF-beta receptor signaling activates SMADs | $3.834\times10^{-6}$ |
| 19 | REAC | REAC:R-HSA-46... | Degradation of DVL | $2.260\times10^{-5}$ |
| 20 | REAC | REAC:R-HSA-17... | Signaling by TGF-beta Receptor Complex | $1.304\times10^{-4}$ |
| 21 | REAC | REAC:R-HSA-21... | Regulation of activated PAK-2p34 by proteasome ... | $1.609\times10^{-4}$ |
| 22 | REAC | REAC:R-HSA-12... | Downregulation of ERBB4 signaling | $2.027\times10^{-4}$ |
| 23 | REAC | REAC:R-HSA-34... | Autodegradation of the E3 ubiquitin ligase COP1 | $2.121\times10^{-4}$ |
| 24 | REAC | REAC:R-HSA-75... | Ubiquitin-dependent degradation of Cyclin D | $2.121\times10^{-4}$ |
| 25 | REAC | REAC:R-HSA-69... | Ubiquitin Mediated Degradation of Phosphorylate... | $2.121\times10^{-4}$ |
| 26 | REAC | REAC:R-HSA-89... | Regulation of RUNX3 expression and activity | $2.764\times10^{-4}$ |
| 27 | REAC | REAC:R-HSA-46... | Degradation of AXIN | $3.143\times10^{-4}$ |
| 28 | REAC | REAC:R-HSA-17... | SCF-beta-TrCP mediated degradation of Emi1 | $3.143\times10^{-4}$ |
| 29 | REAC | REAC:R-HSA-88... | PTK6 Regulates RTKs and Their Effectors AKT1 and... | $3.623\times10^{-4}$ |
| 30 | REAC | REAC:R-HSA-90... | Signaling by NOTCH4 | $3.643\times10^{-4}$ |
| 31 | REAC | REAC:R-HSA-56... | NIK-->noncanonical NF-kB signaling | $5.121\times10^{-4}$ |
| 32 | REAC | REAC:R-HSA-56... | Hedgehog 'on' state | $5.281\times10^{-4}$ |
| 33 | REAC | REAC:R-HSA-18... | SCF(Skp2)-mediated degradation of p27/p21 | $5.752\times10^{-4}$ |
| 34 | REAC | REAC:R-HSA-56... | GLI3 is processed to GLI3R by the proteasome | $5.752\times10^{-4}$ |
| 35 | REAC | REAC:R-HSA-56... | Degradation of GLI2 by the proteasome | $5.752\times10^{-4}$ |
| 36 | REAC | REAC:R-HSA-56... | Degradation of GLI1 by the proteasome | $5.752\times10^{-4}$ |
| 37 | REAC | REAC:R-HSA-93... | IRAK2 mediated activation of TAK1 complex | $5.993\times10^{-4}$ |
| 38 | REAC | REAC:R-HSA-56... | Dectin-1 mediated noncanonical NF-kB signaling | $5.752\times10^{-4}$ |
| 39 | REAC | REAC:R-HSA-90... | TICAM1,TRAF6-dependent induction of TAK1 com... | $9.349\times10^{-4}$ |
| 40 | REAC | REAC:R-HSA-53... | Hedgehog ligand biogenesis | $9.971\times10^{-4}$ |
| 41 | REAC | REAC:R-HSA-17... | APC/C:Cdc20 mediated degradation of Securin | $1.356\times10^{-3}$ |
| 42 | REAC | REAC:R-HSA-13... | Downregulation of ERBB2:ERBB3 signaling | $1.392\times10^{-3}$ |
| 43 | REAC | REAC:R-HSA-90... | TICAM1-dependent activation of IRF3/IRF7 | $1.392\times10^{-3}$ |
| 44 | REAC | REAC:R-HSA-90... | Signaling by TGFB family members | $1.815\times10^{-3}$ |
| 45 | REAC | REAC:R-HSA-17... | Cdc20:Phospho-APC/C mediated degradation of C... | $2.192\times10^{-3}$ |
| 46 | REAC | REAC:R-HSA-17... | APC/C:Cdh1 mediated degradation of Cdc20 and ... | $2.402\times10^{-3}$ |
| 47 | REAC | REAC:R-HSA-18... | EGFR downregulation | $3.235\times10^{-3}$ |
| 48 | REAC | REAC:R-HSA-97... | IRAK1 recruits IKK complex upon TLR7/8 or 9 stim... | $2.774\times10^{-3}$ |
| 49 | WP | WP:WP183 | Proteasome Degradation | $4.742\times10^{-2}$ |
| 50 | TF | TF:M04515_1 | Factor: E2F-1; motif: WWTGGCGCCAAA; match cla... | $1.641\times10^{-5}$ |
| 51 | TF | TF:M04826_1 | Factor: p300; motif: ACNTCCG; match class: 1 | $1.561\times10^{-4}$ |
| 52 | TF | TF:M11530 | Factor: E2F-2; motif: NWTTTGGCGCCAWWNN | $1.912\times10^{-4}$ |
| 53 | TF | TF:M10438 | Factor: ZF5; motif: GGSGCGCGS | $5.816\times10^{-4}$ |
| 54 | TF | TF:M00932_1 | Factor: Sp1; motif: NNGGGGCGGGGNN; match cla... | $8.032\times10^{-4}$ |
| 55 | TF | TF:M12160 | Factor: KLF15; motif: RCCMCRCCCMCN | $1.247\times10^{-2}$ |
| 56 | TF | TF:M03925 | Factor: YY2; motif: NCCGCCATNTY | $3.091\times10^{-2}$ |
| 57 | TF | TF:M03924_1 | Factor: YY1; motif: NNCGCCATTNN; match class: 1 | $4.267\times10^{-2}$ |
| 58 | TF | TF:M10530 | Factor: sp4; motif: NNNGCYCCGCCCCCY | $3.445\times10^{-2}$ |
| 59 | MIRNA | MIRNA:hsa-miR... | hsa-miR-615-3p | $5.288\times10^{-11}$ |
| 60 | MIRNA | MIRNA:hsa-miR... | hsa-miR-324-3p | $2.115\times10^{-2}$ |

| | |
|---|---|
| **version** | e103_eg50_p15_68c0e33 |
| **date** | 5/18/2021, 10:26:39 PM |
| **organism** | hsapiens |

g:Profiler

**Supplementary Table 3: GO terms list for the Manhattan plot (Figure 3A) provided from g:Profiler software for the significantly downregulated proteins on statin treatment in whole lysate proteomics MS-MS analysis**. The color coding is dependent on the significance value (p value) assigned to the terms in the software based on the protein list belonging to the term and the statistical significance of their match, blue signifying highly significant to orange signifying least significant. Therefore, all the GO terms listed are highly significant.

| S.No. | Gene name | qPCR Primer sequence |
|---|---|---|
| 1 | SATB1 | Forward- AACTCAGGGCTTGCTTTC<br>Reverse- CCTGGTATATTCGGTCTCTTTC |
| 2 | SATB2 | Forward- AGAGATGAACCAGAGCACATTAG<br>Reverse- GTTGCTGACACATTGGCATAAT |
| 3 | β-catenin | Forward- AAGGTGTGGCGACATATGCA<br>Reverse- GTAATCTTGTGGCTTGTCCTCAGA |
| 4 | 18s rRNA | Forward- CGCCGCTAGAGGTGAAATTCT<br>Reverse- CGAACCTCCGACTTTCGTTCT |
| 5 | SREBF2 | Forward- GCTGCCAAGGAGAGTCTAT<br>Reverse- AGGTTTCACCAAGGACTCTAT |
| 6 | HMGR | Forward- ACAAGAATTTAGTGGGCTCTG<br>Reverse- TCCTGTCCACAGGCAAT |
| 7 | CDH1 | Forward- GGCTGGACCGAGAGAGTTTC<br>Reverse- CCTGACCCTTGTACGTGGTG |
| 8 | Vimentin | Forward- GCGAGGAGAGCAGGATTTCT<br>Reverse- TGGGTATCAACCAGAGGGAGT |
| 9 | E-cadherin | Forward- GTCCTGGGCAGAGTGAAT<br>Reverse- GGGTTATGAAACCGTAGAGGC |

**Supplementary Table 4: The list of qPCR primers used for the analysis.**

| Patient # | Pre-treatment | | Post-treatment | |
|---|---|---|---|---|
| | SATB1 score | SATB2 score | SATB1 score | SATB2 score |
| 1 | 1 | 2 | 1 | 1 |
| 2 | 0 | 0 | 0 | 0 |
| 3 | 1 | 0 | 1 | 2 |
| 4 | 1 | 2 | 0 | 6 |
| 5 | 0 | 0 | 0 | 3 |
| 6 | 4 | 9 | 0 | 0 |
| 7 | 2 | 1 | 0 | 0 |
| 8 | 0 | 6 | 0 | 1 |
| 9 | 1 | 0 | 0 | 1 |
| 10 | 0 | 0 | 0 | 0 |

**Supplementary Table 5: Immunohistological H scores for SATB1 and SATB2 signal intensities of paired biopsy samples before and after statin treatment.** Calculation of the H score is based on signal intensity and percentage of cells positive for signal. The scoring for the two parameters is then multiplied to assign a final H score. In terms of the number of cells showing signal, 1 = less than 33% cells positive, 2 = 33-66% cells positive, 3 = more than 66% cells positive. In terms of intensity 1 = weak, 2 = moderate, 3 = high intensity. H Scores for 10 patients are tabulated here.

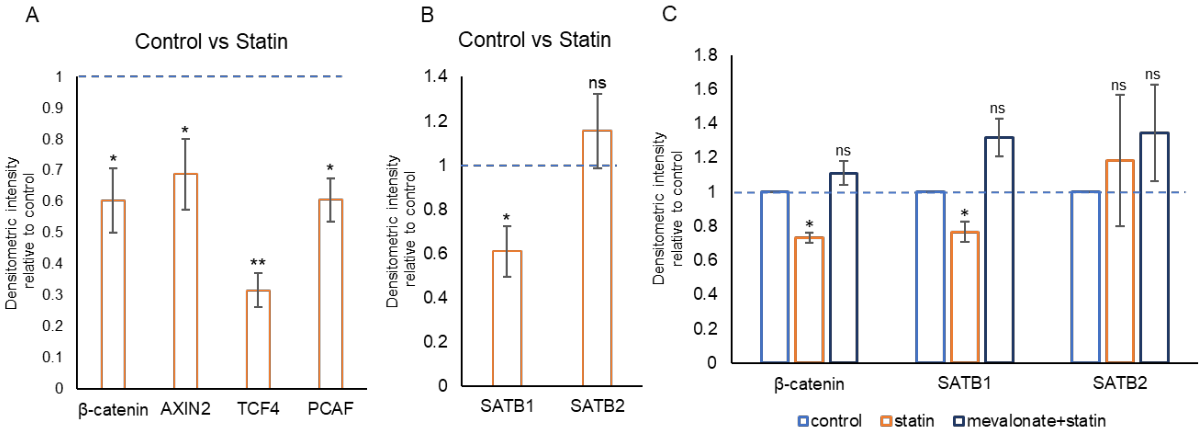


**Supplementary Figure 1: Densitometric expression profiling of Wnt pathway proteins in statin-treated cells as compared to control.** (A) Densitometric intensity relative to control for immunoblots of β-catenin, AXIN2, TCF4 and PCAF depicted in Figure 3B. All proteins exhibited a significant reduction upon statin treatment. (B) Densitometric intensity relative to control for immunoblots of SATB1 and SATB2 in Figure 3D. SATB1 is observed to be significantly downregulated, whereas SATB2 did not exhibit a significant alteration. (C) Densitometric analysis relative to control for immunoblots of β-catenin, SATB1 and SATB2 in Figure 4A. Statin treatment resulted in a reduction in protein levels of β-catenin and SATB1, whereas mevalonate plus statin treated cells exhibit a rescue. However, SATB2 levels were not significantly altered in both statin and mevalonate plus statin treated cells (Biological replicates n=3, *$p<0.05$, **$p<0.005$, ns stands for non-significant upon Students' t test analysis).

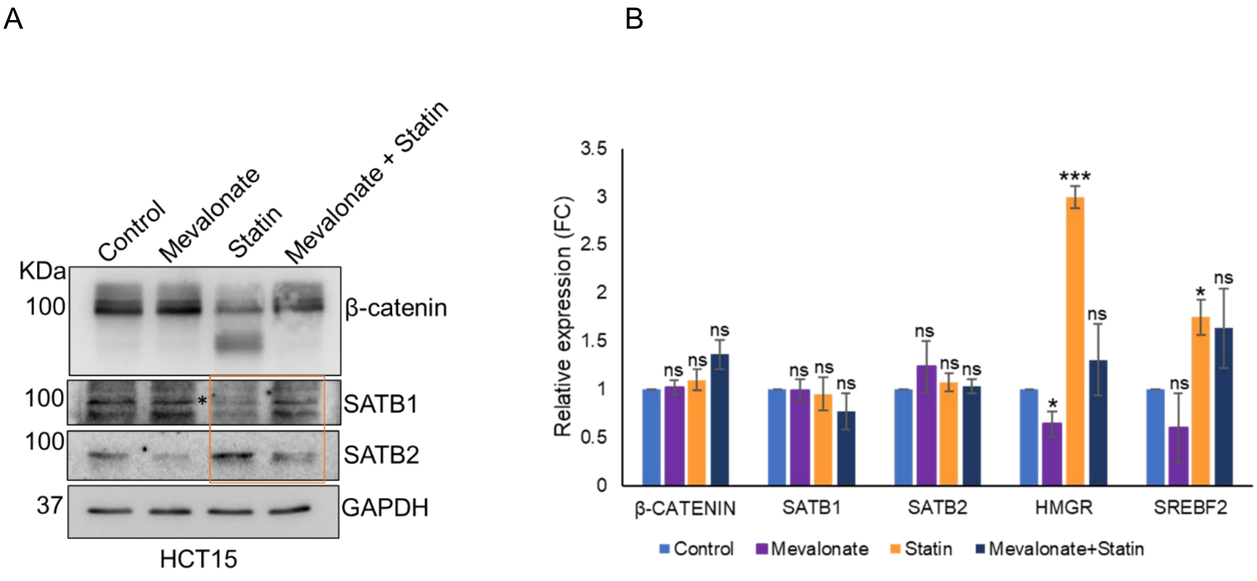


**Supplementary Figure 2: Mevalonate supplementation along with statin treatment rescues the protein levels of SATB1.** (A) Immunoblot for β-catenin, SATB1 and SATB2 protein expression in mevalonate, mevalonate + statin treatment conditions. The upper band (* outside the red box) in the SATB1 blot denotes the band corresponding to its expected mobility in SDS-PAGE. The protein level reduced post statin treatment is restored upon mevalonate supplement. SATB2 levels remain comparable to the untreated control on addition of mevalonate. Statin treatment resulted in slight upregulation of SATB2 protein, however, is restored to the level observed in untreated set. (B) Relative gene expression of β-catenin, SATB1, SATB2, HMGR and SREBF2 respectively in statin, mevalonate and mevalonate + statin conditions. No alteration in transcript levels of β-catenin, SATB1 and SATB2 was observed. The cholesterol responsive genes HMGR and SREBF2 exhibit upregulation upon statin treatment, whereas the levels are restored upon mevalonate plus statin treatment (Biological replicates n=3, *$p<0.05$, ***$p<0.0005$, ns stands for non-significant by Students' t test analysis).

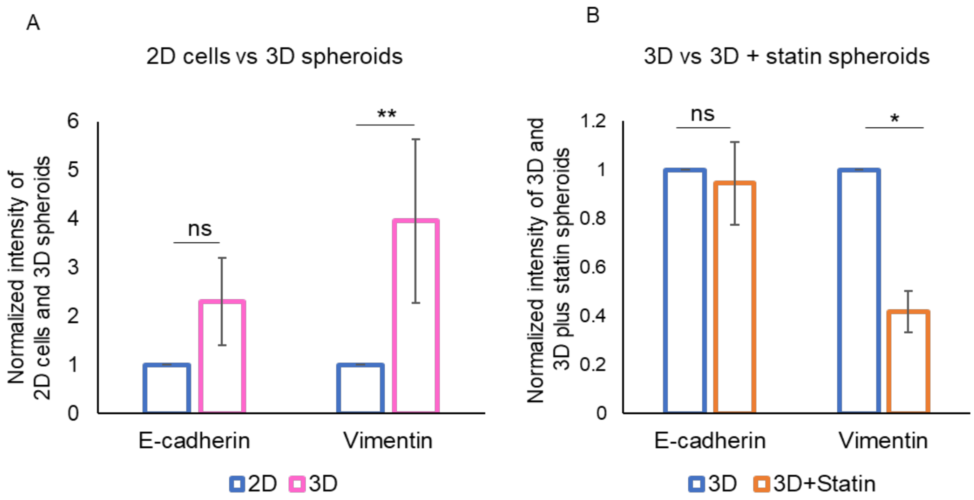


**Supplementary Figure 3: Analysis of EMT markers in 2D cells versus 3D spheroids and vehicle control treated spheroids versus statin-treated spheroids.** (A) Densitometric analysis of intensity relative to 2D cells for immunoblots of E-cadherin and Vimentin in 3D spheroids depicted in Figure 5C. E-cadherin levels were not altered significantly in 2D cells and 3D spheroids. However, Vimentin levels were significantly upregulated in spheroids, suggesting an EMT phenotype. (B) Densitometric analysis of the normalized intensities of immunoblots for E-cadherin and Vimentin upon statin treatment in spheroids depicted in Figure 6D. Upon statin treatment the spheroids exhibited significantly reduced Vimentin level suggesting a reversal of the EMT phenotype. However, E-cadherin levels were not significantly altered (Biological replicates n=3, *p<0.05, **p<0.005, ns stands for non-significant upon Students' t test analysis).

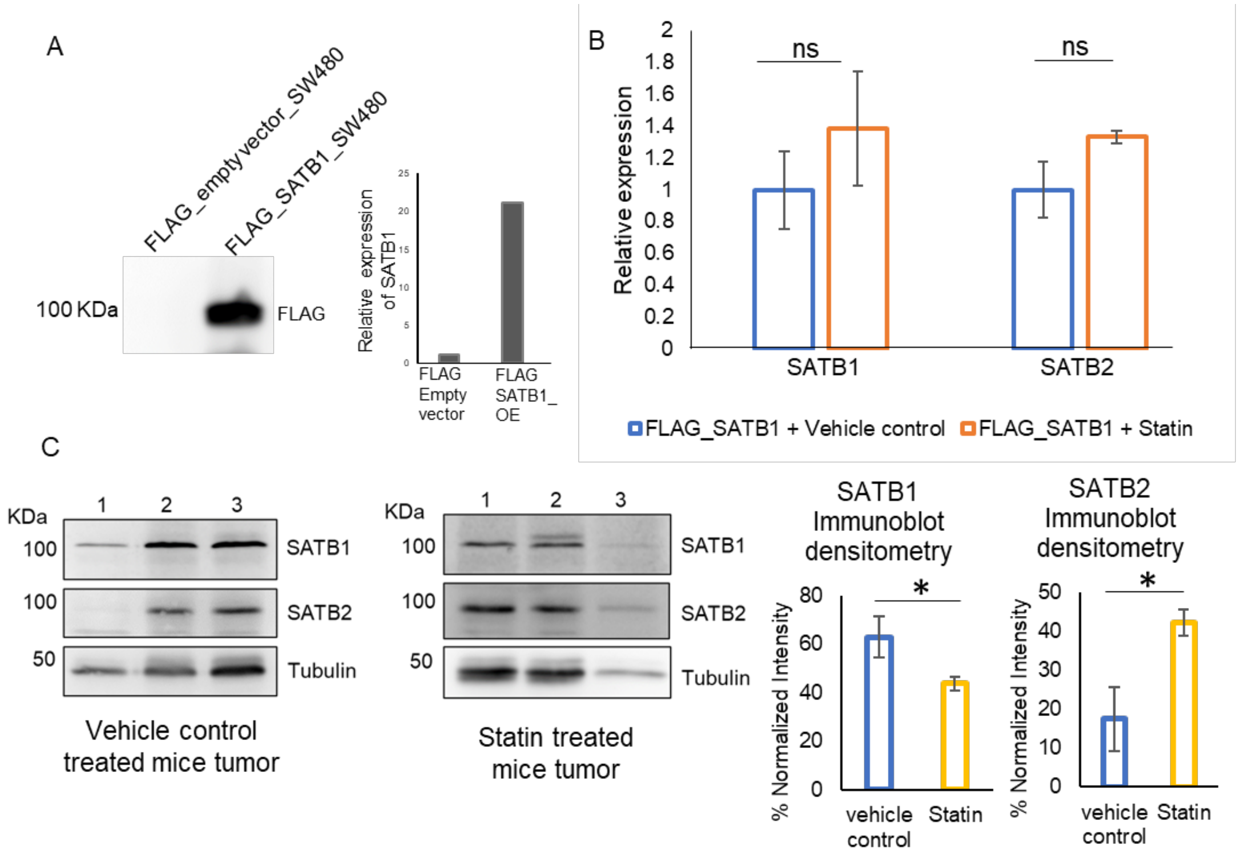


**Supplementary Figure 4: Molecular profiling of the tumor tissues isolated from statin treated NOD-SCID mice suggests downregulation of SATB1.** (A) Validation of over-expression of FLAG-SATB1 in SW480 cell line injected in mice, at both protein as well as transcript level. (B) Transcript levels of SATB1 and SATB2 in tumor tissues harvested from mice on vehicle control treatment and statin treatment respectively, show no alteration. (C) Immunoblots for SATB1 and SATB2 protein levels in tumors obtained from vehicle control and statin treated sets of mice. SATB1 levels seem to be reduced on statin treatment, corroborating the *in cellulo* data. The densitometry analysis graph on the right demonstrates a significant downregulation of SATB1 in statin treated mice tumor samples (Biological replicates n=3, *p<0.05 and ns stands for non-significant by Students' T test).

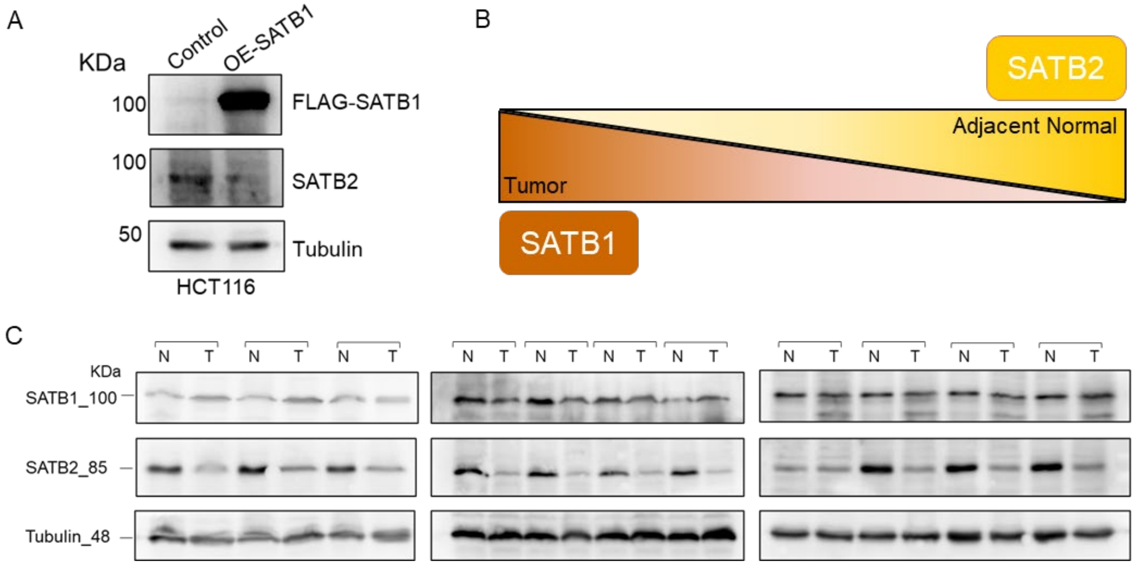


**Supplementary Figure 5: Signature reciprocal expression profile of SATB proteins.** (A) Ectopic over-expression of SATB1 in HCT116 cells demonstrated a reciprocal effect on SATB2 expression at protein level. As SATB1 expression is increased SATB2 expression is significantly reduced at protein level. (B) Schematic representation of the contrasting expression profile of SATB proteins in tumor biopsies and adjacent normal tissues. SATB1 is higher in tumor tissue as compared to SATB2 expression which is higher in adjacent normal. (C) Validation of the reciprocal expression profile of SATB proteins in tumor and normal adjacent biopsy tissue samples. A total of 11 patients, with paired samples of adjacent normal (N) and tumor tissue (T) were probed for SATB1 and SATB2 expression by immunoblotting using respective antibodies. The tumor samples exhibit higher expression of SATB1 than SATB2. In contrast, SATB2 expression is higher in the adjacent normal tissue samples. These patient biopsies were obtained during regular checkups for adenoma prognosis as part of the human trial conducted at Tata Memorial Hospital, Mumbai, India.

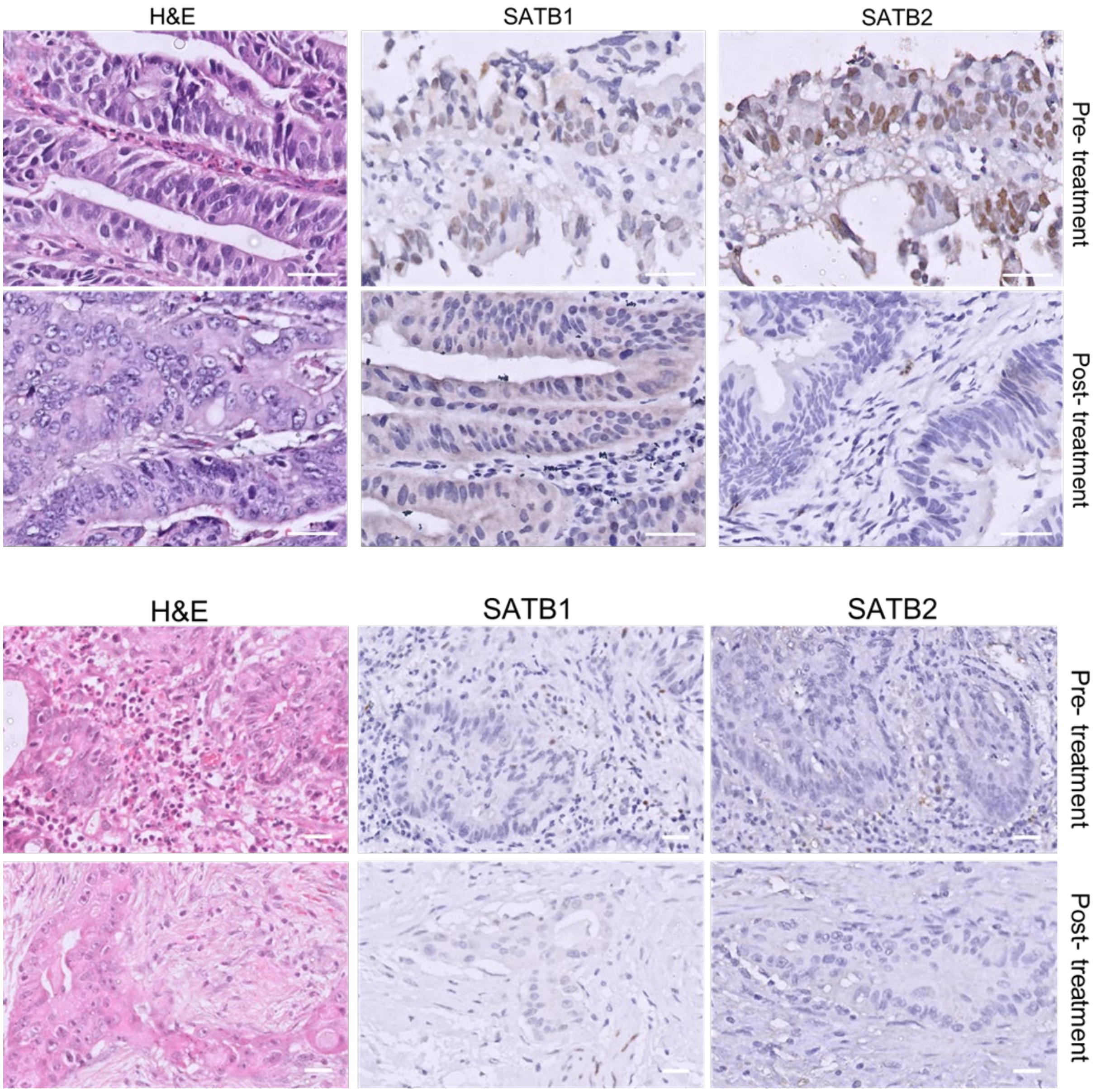


**Supplementary Figure 6: Representative images of immunohistochemistry staining for tissue biopsies.** The sections of biopsies were obtained from two different patients in arm ‘A’, in pre-statin treatment and post-treatment conditions (scale bar 350 μm has been kept constant). The intensity of staining for SATB1 and SATB2 proteins was quantified using the H score. The adenoma tissues in both the panels exhibit a reduction in the expression of SATB1 in post-statin treated as compared to the pre-treatment samples. In comparison, SATB2 expression seems variable.

**Uncropped immunoblot data**

Figure 3B:

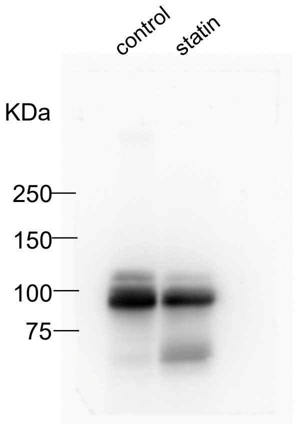


IB: β-catenin

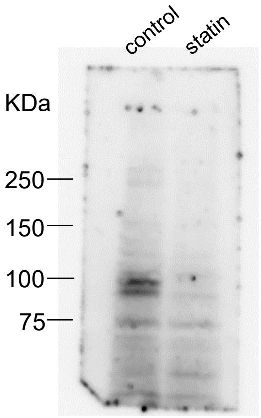


IB: AXIN2

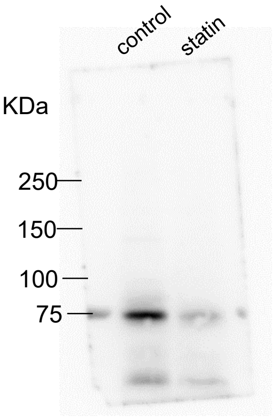


IB: TCF4

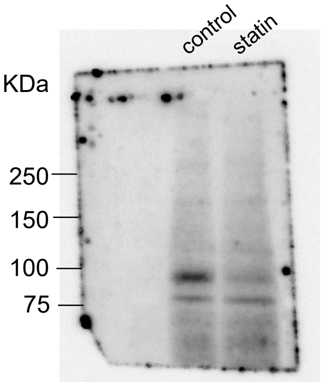


IB: PCAF

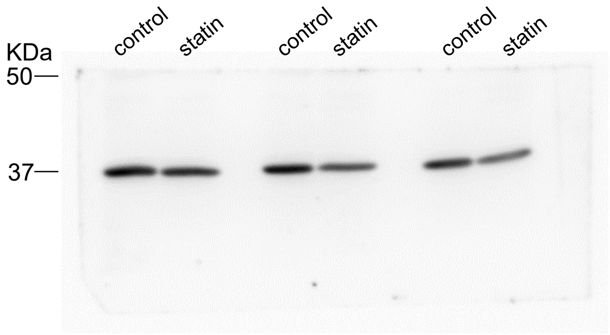


IB: GAPDH

Figure 3D:

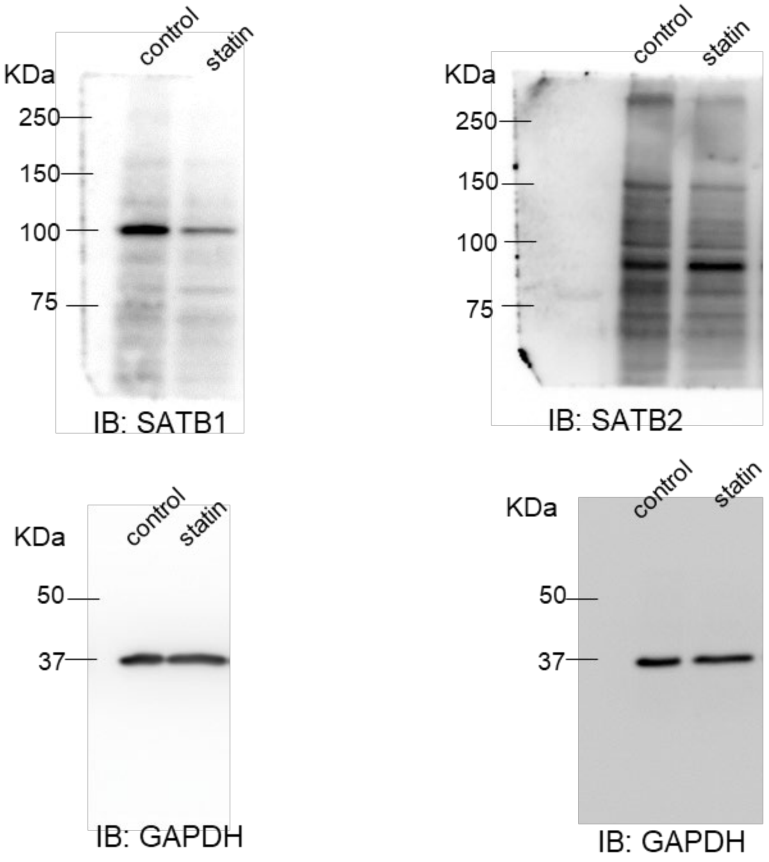
control
statin
KDa
250
150
100
75
IB: SATB1
control
statin
KDa
250
150
100
75
IB: SATB2
control
statin
KDa
50
37
IB: GAPDH
control
statin
KDa
50
37
IB: GAPDH

Figure 4A:

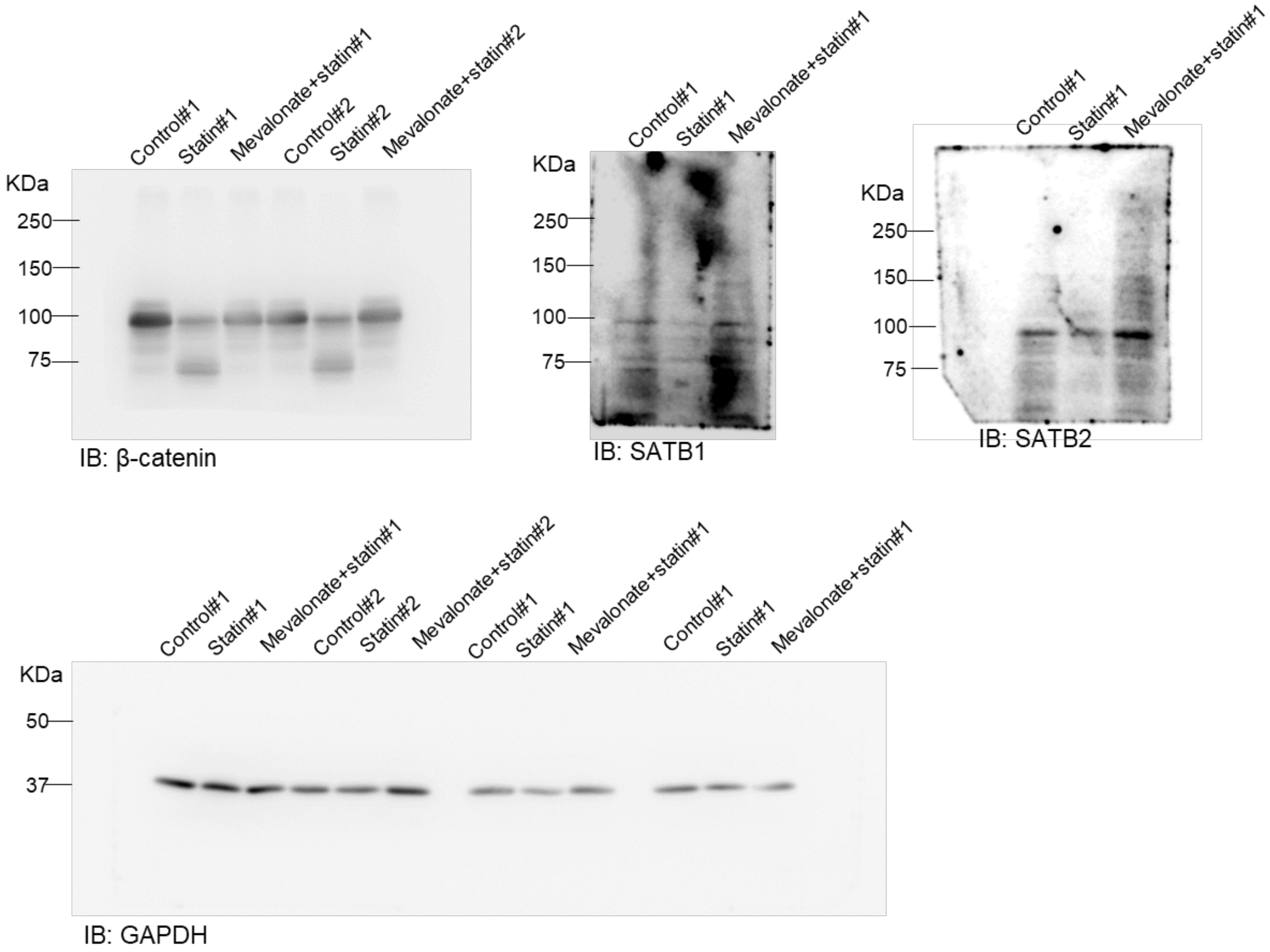
Control#1
Statin#1
Mevalonate+statin#1
Control#2
Statin#2
Mevalonate+statin#2
KDa
250
150
100
75
IB: β-catenin
Control#1
Statin#1
Mevalonate+statin#1
KDa
250
150
100
75
IB: SATB1
Control#1
Statin#1
Mevalonate+statin#1
KDa
250
150
100
75
IB: SATB2
Control#1
Statin#1
Mevalonate+statin#1
Control#2
Statin#2
Mevalonate+statin#2
Control#1
Statin#1
Mevalonate+statin#1
Control#1
Statin#1
Mevalonate+statin#1
KDa
50
37
IB: GAPDH

Figure 5C:

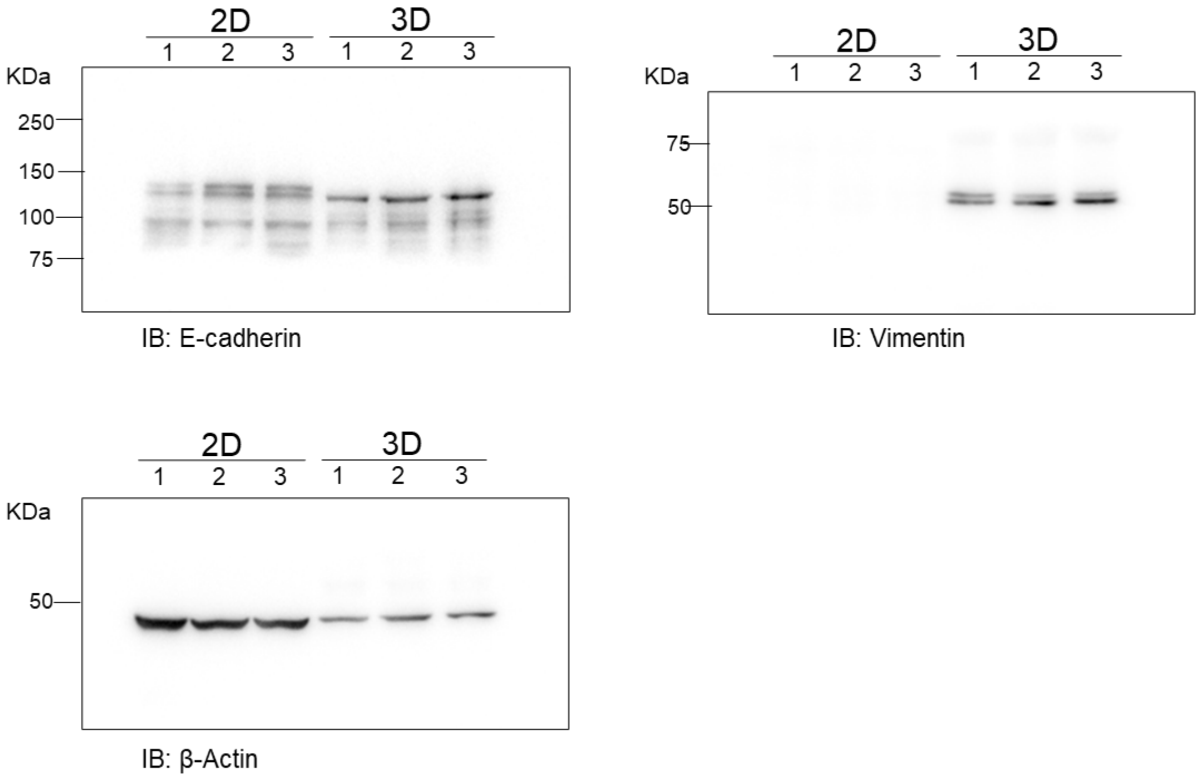


Figure 5D:

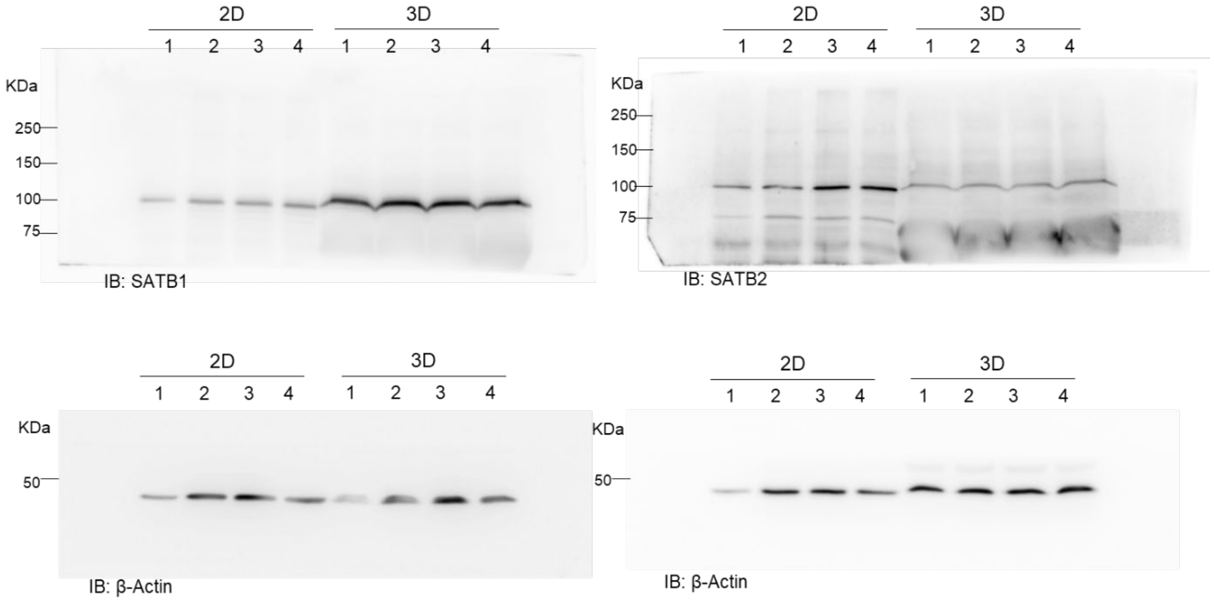

Figure 6D:

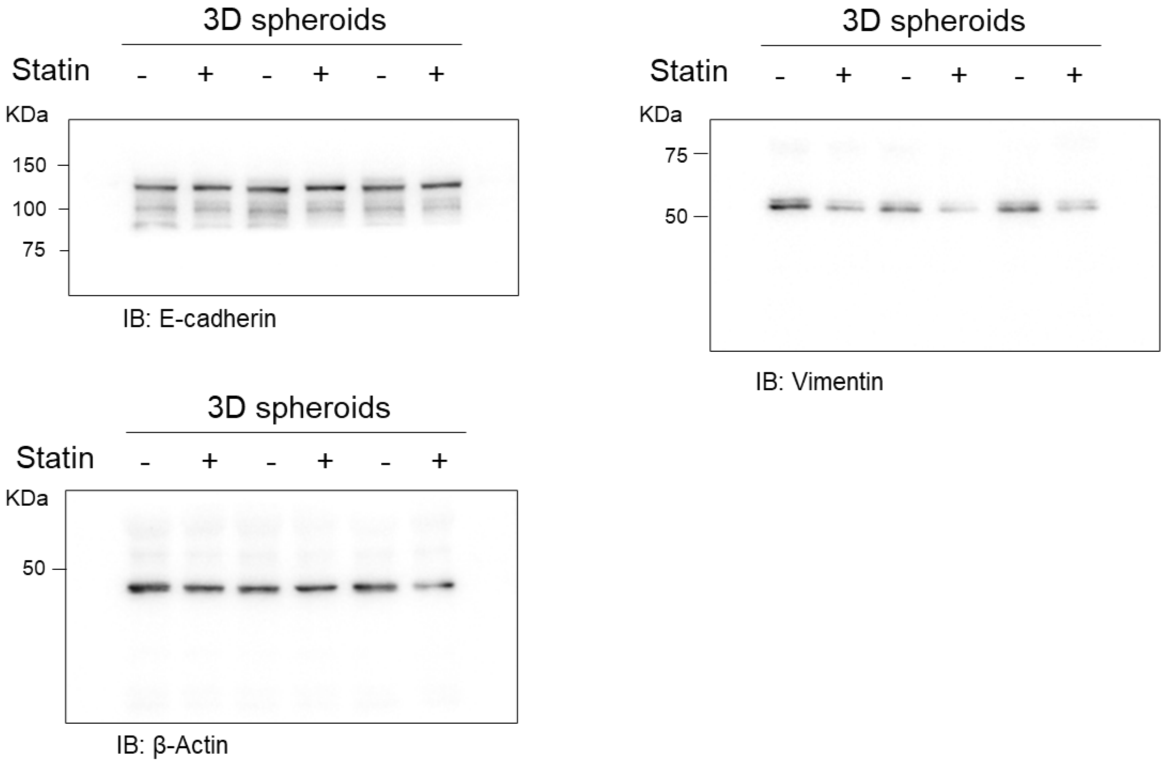


Figure 6E:

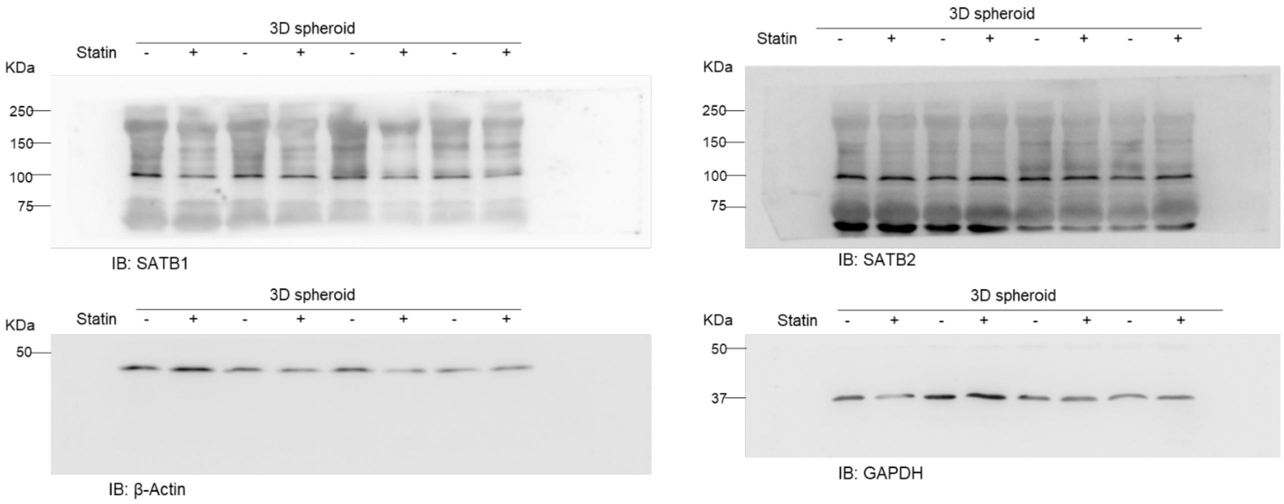

Supplementary Figure 2A:

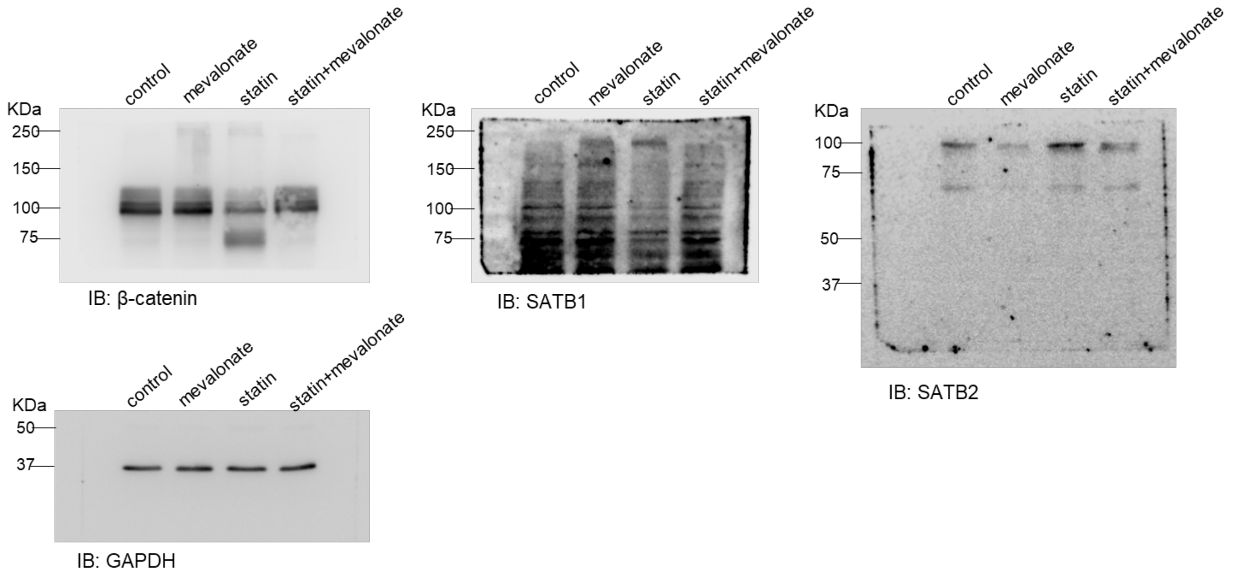


Supplementary Figure 4C:

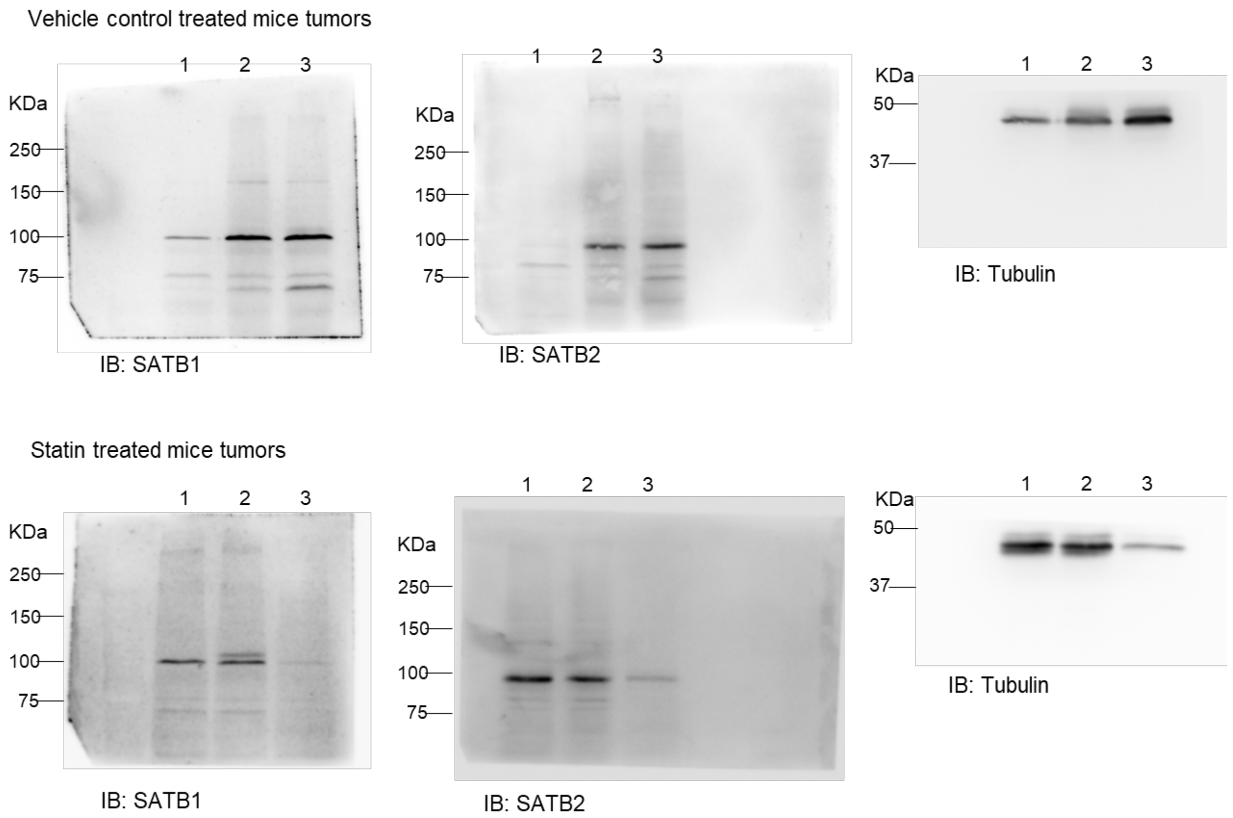

Supplementary Figure 5A:

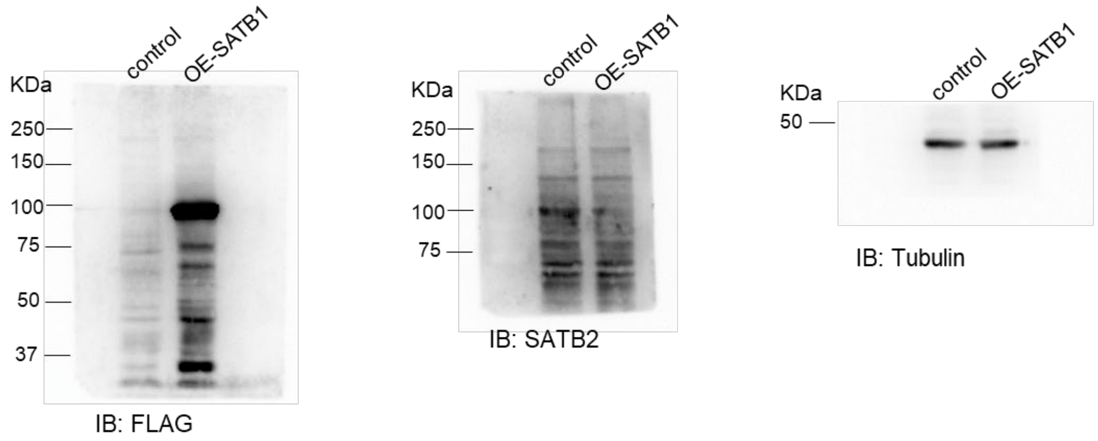


Supplementary Figure 5C:

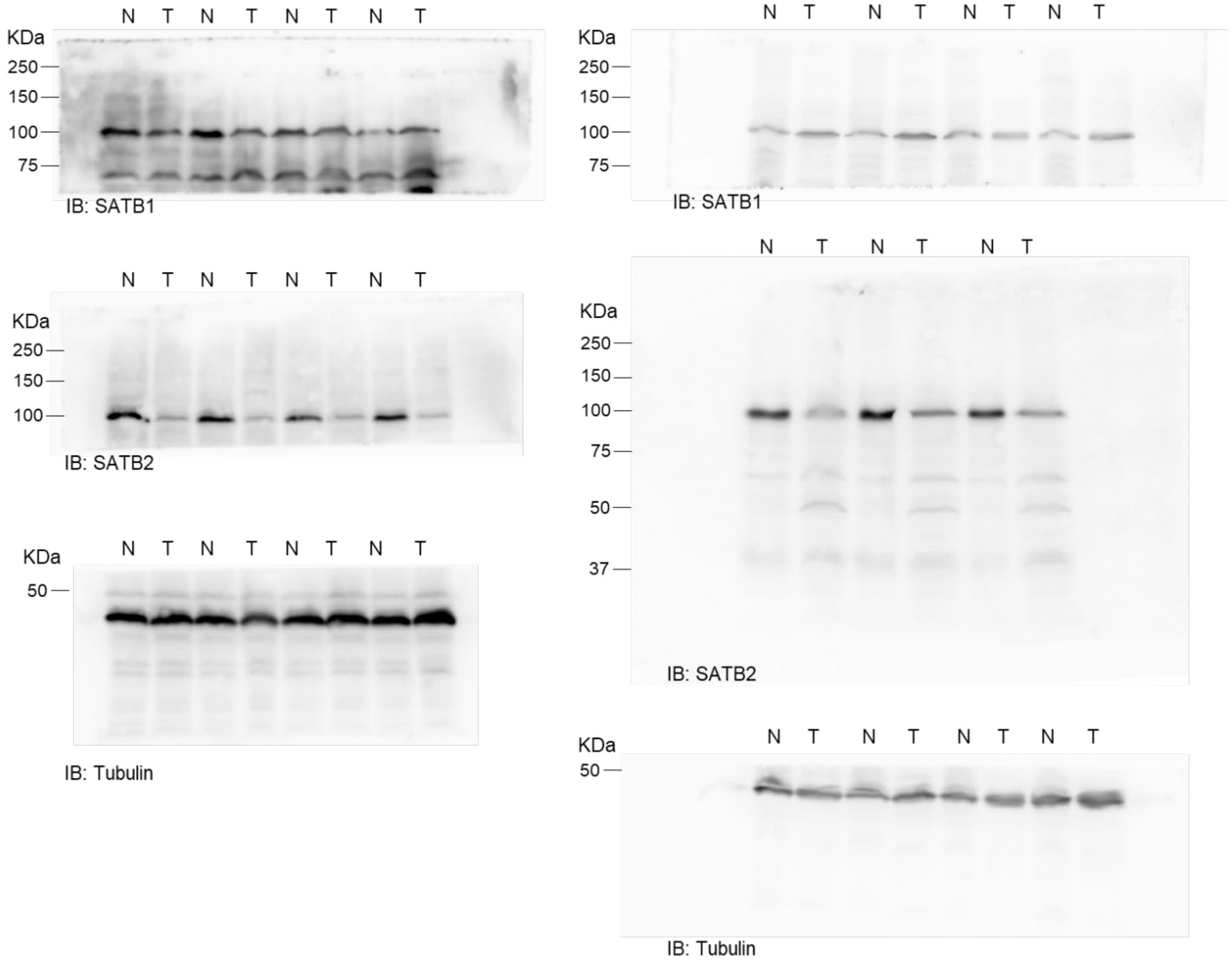

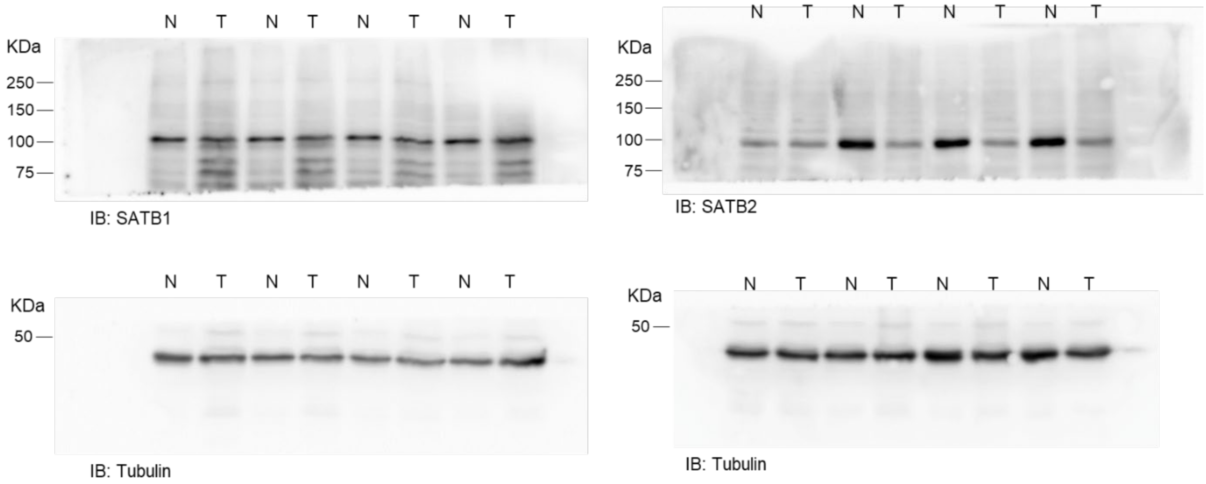
N T N T N T N T
KDa
250
150
100
75
IB: SATB1
N T N T N T N T
KDa
250
150
100
75
IB: SATB2
N T N T N T N T
KDa
50
IB: Tubulin
N T N T N T N T
KDa
50
IB: Tubulin
